# Untargeted metabolomics reveals anion and organ-specific biochemistry of salinity tolerance in willow

**DOI:** 10.1101/2024.07.16.603755

**Authors:** Eszter Sas, Adrien Frémont, Emmanuel Gonzalez, Mathieu Sarrazin, Simon Barnabé, Michel Labrecque, Nicholas James Beresford Brereton, Frédéric Emmanuel Pitre

## Abstract

Willows can alleviate soil salinisation while generating sustainable feedstock for biorefinery, yet the metabolomic adaptations underlying their salt tolerance remain poorly understood. Testing two environmentally abundant salts, the response of *Salix miyabeana* was assessed after treatment with a moderate concentration of NaCl, and both moderate and high concentrations of Na_2_SO_4_ in a 12-week pot trial. Willows tolerated salts across all treatments (up to 9.1dS m^-1^ soil EC_e_), maintaining photosynthesis and biomass while selectively partitioning ions, confining Na^+^ to roots and accumulating Cl^-^ and SO_4_ ^2-^ in the canopy, and adapting to osmotic stress via reduced stomatal conductance. Untargeted LC-MS/MS captured over 5,000 putative compounds, characterising the baseline willow metabolome, including 278 core compounds constitutively produced across organs. Comparative statistical analyses revealed widespread metabolic reprogramming in response to soil salinity, altering 28% of the overall metabolome, and highlighting organ-tailored regulation. Comparing both salt forms at equimolar sodium, generalised salinity responses were limited to 3% of the metabolome, predominantly in roots. Anion-specific metabolomic responses were more extensive, with NaCl reducing carbohydrates and TCA intermediates, thereby exerting pressure on carbon and energy resources, alongside the accumulation of root structuring compounds, antioxidants flavonoids, and fatty acids. In contrast, Na_2_SO_4_ salinity triggered accumulation of sulphur-containing larger peptides, suggesting that excess sulphate incorporation leverage ion toxicity to produce specialized salt-tolerance associated metabolites. This high-depth picture of the willow metabolome underscores the importance of capturing plant adaptations to salt stress at organ-scale and considering ion-specific contributions to soil salinity.

## 2 Introduction

Soil salinisation represents a major threat to agricultural productivity and environmental health on a global scale. It is estimated that over 20% of irrigated cropland is impacted by salt accumulation (FAO, 2015), reducing yields and critical ecosystem biodiversity (Zörb et al., 2019). Climate change and human activities are predicted to intensify both natural and anthropogenic soil salinisation processes, further threatening food security (Singh, 2021). While salt stress research typically investigates NaCl, Na_2_SO_4_, derived from anthropogenic sources like industrial sulphur emissions and agricultural practices, as well as natural volcanic and marine environments (Dick et al., 2008), represents another widespread salt. The use of salt-tolerant biomass crops able to sequester salt and improve soil health, could help reclaim and repurpose salt-impacted areas (Abreu et al., 2022; Quinn et al., 2015).

Soil salinisation results from soluble ions accumulation such as Na^+^, K^+^, Mg^2+^, Ca^2+^, Cl^-^, SO_4_^2-^ and CO_3_^2-^. When in contact with roots, these ions can induce both osmotic stress and ion toxicity (Munns and Tester, 2008). The osmotic component, primarily driven by any dissolved salt concentration, alters water potential gradients, disrupting cell turgor pressure and elongation. In response, stomatal closure limits water loss via transpiration but also impairs photosynthate supply, ultimately constraining growth through both water deficit and reduced carbon availability (Munns et al., 2020b). Ion toxicity, however, is dependent on the nature of the ion, with different elements inducing unique patterns of signalling, transport, oxidative damage, energy allocation, and hormone-regulated growth processes (Llanes et al., 2013; Reginato et al., 2021; Richter et al., 2019).

Chloride (Cl^-^), an essential micronutrient for plants, contributes to vital physiological processes (Wege et al., 2017). As a major osmotically active solute, chloride fluxes are involved in vacuolar turgor required for cell expansion and are implicated in stomatal aperture regulation, contributing to water balance and gas exchange (Teakle and Tyerman, 2010). Additionally, chloride acts as a counter-ion, regulating membrane potential, intracellular pH gradients, electrical signalling in the cytoplasm (White and Broadley, 2001) and serves as a cofactor in the oxygen-evolving complex of photosystem II (Ifuku, 2015). However, in salt-impacted soils, excess chloride damages plants, leading to impaired photosynthesis, reduced NO_3_^-^ uptake (Abdelgadir et al., 2005) and general enzymatic activity (Geilfus, 2018). In contrast, sulphate (SO_4_^2-^), the major bioavailable form of sulphur, is an essential macronutrient involved in physiological and biochemical processes (Prasad and Shivay, 2018; Watanabe and Hoefgen, 2019). Sulphur is required for the production of the amino acids cysteine and methionine, peptide derivatives such as glutathione, and disulphide bonds contributing to enzyme activity and stability. The production of chlorophyll, vitamins and certain plant defence specialised metabolites, such as glucosinolates, also depend upon sulphur (Chan et al., 2019; Rausch and Wachter, 2005). Plants regulate sulphate uptake and assimilation through sulphate specific transporters to avoid toxicity (Buchner et al., 2004), however in some crop species, such as *Brassica rapa*, excess sulphate ions can create nutrient deficiency by competing with the absorption of other ions, especially calcium and phosphorus (Reich et al., 2017).

Plants mediate salt stress through a coordinated series of physiological, structural, biochemical and molecular adjustments (Arif et al., 2020; Deinlein et al., 2014; Gupta and Huang, 2014; van Zelm et al., 2020; Yang and Guo, 2018). Although pathways vary between plant species (Sanchez et al., 2008) and depend on the type of ions involved (Reich et al., 2017; Richter et al., 2019), key tolerance mechanisms include osmotic regulation, ion homeostasis, antioxidant defence, and growth modulation. A primary adaptive response to osmotic stress is the accumulation of osmolytes, which regulate water status by reducing cell osmotic potential (D’Amelia et al., 2018). This common osmotic stress response is not exclusively induced by excess salt nor is dependent on salt type (Munns, 2002). Osmolytes include inorganic ions (Rodriguez et al., 1997; Wege et al., 2017) and organic compounds such as amino acids (e.g. proline), polyamines (e.g. spermidine), quaternary ammonium compounds (e.g. glycine betaine), sugars (e.g. sucrose, trehalose), and polyols (e.g. mannitol, sorbitol) (D’Amelia et al., 2018; Slama et al., 2015; Yang and Guo, 2018). While the biosynthesis of these osmoprotectants imposes a metabolic energy cost (Munns et al., 2020a), their functions extend beyond osmotic adjustment to detoxification processes such as antioxidant activity and chaperone-mediated protein stabilisation (Yancey, 2005). The upregulation of enzymatic (e.g. superoxide dismutase SOD, catalase CAT, peroxidase POX) and non-enzymatic (e.g. glutathione, ascorbate, flavonoids, anthocyanins) antioxidants is crucial for mitigating damage to cellular components such as lipids, proteins, and DNA caused by reactive oxygen species (ROS) (Gill and Tuteja, 2010a). More generally, plants avoid cytotoxicity and maintain ion homoeostasis through salt exclusion from roots, compartmentalisation into the vacuole or apoplast, selective transport and excretion through salt glands. Additionally, hormonal signalling (Ryu and Cho, 2015), modification of root architecture (Zou et al., 2022), xylem alteration and loading (Janz et al., 2012) and symbiotic associations with microorganisms such as arbuscular mycorrhizal fungi (Evelin et al., 2009) also play significant roles in enhancing plant tolerance to salinity.

Salicaceae, including poplars (*Populus sp.)* and willows (*Salix sp.*), are common phytoremediation and biorefinery crops on marginal lands (Brereton et al., 2017; Dimitriou and Aronsson, 2005; Gonzalez et al., 2018; Rodzkin and Volk, 2018; Sas et al., 2021; Volk et al., 2006), with high potential for growth on salinised soils. Some cultivars can maintain transpiration and growth under saline conditions up to 6-8 dS m^-1^ (Bilek et al., 2020; Chen and Polle, 2010; Huang et al., 2020; Mirck and Zalesny, 2015). This tolerance likely engages high metabolic diversity, including reactive oxygen species (ROS) protective phenolics and other antioxidants (Sui and Wang, 2020; Zhou et al., 2020). However, understanding of the metabolomic toolkit employed beyond these compounds, and their variation across different organs and salts, remains limited.

In this study, the shared and unique metabolomic adjustments to moderate NaCl, moderate Na_2_SO_4_ and high Na_2_SO_4_ salinity treatment are assessed across multiple organs (Fig.1) to determine generalised and ion-specific salt tolerance metabolome in willow.

**Figure 1:**
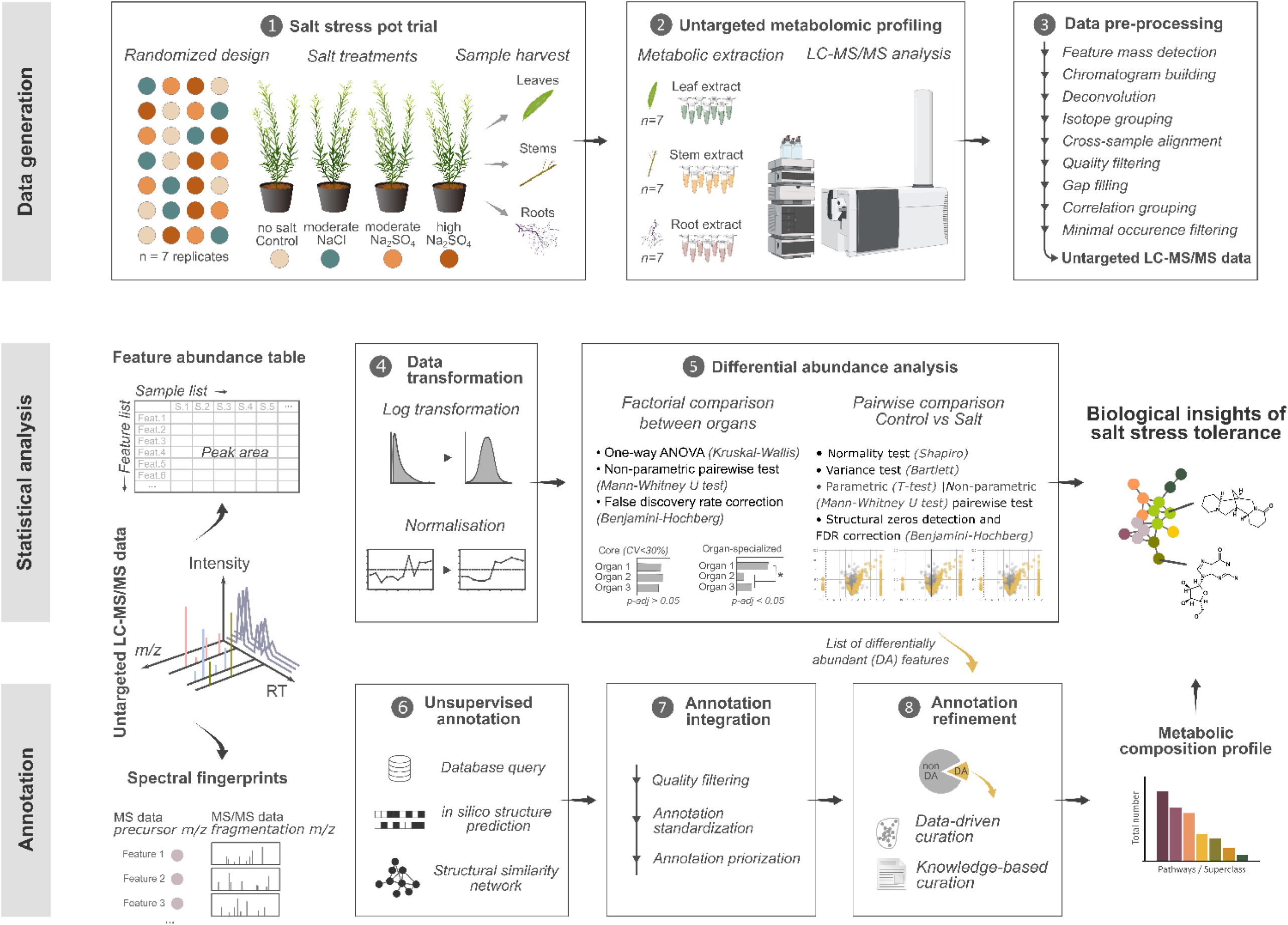
Workflow for untargeted metabolomics approach. A randomized pot trial treated willows with salt solutions or water controls (7 replicates per treatment) to explore variation in anion-specific salt tolerance mechanisms across organs. Metabolites were extracted from root, stem and leaf samples and analysed by LC-MS/MS. Resulting chromatographic data were pre-processed using MZmine for features mass detection, cross-sample alignment and quality filters. Statistical analysis used a cross-sample aligned peak area table. After log transformation and Eigen MS normalization, univariate analysis was used to factorially compare feature peak areas across willow organs in controls, while pairwise comparisons were used to identify differentially abundant (DA) compounds between each salt treatment compared to controls. Parametric or non-parametric tests were used based on feature peak area normality and variance homogeneity before a false discovery rate correction was applied to account for multiple testing. Annotation strategy used precursors information and corresponding spectral fingerprints (when available) to query multiple databases and for *in silico* structure prediction to generate a list of annotation candidates (unsupervised annotation). After standardizing compounds identity information and assigning prioritization criteria, the best candidate was retained. Annotation was further refined for compounds identified as differentially abundant due to salt treatment and used data-driven (structural similarity networking) and knowledge-driven (literature) curation.

## 3 Methods

### 3.1 Experimental design, plant growth, sample collection and physiological traits

The willow cultivar *Salix miyabeana* ‘SX64’ was grown from cuttings in a polyethylene grow tunnel within 15 L pots filled with all-purpose growing soil (Promix^TM^), sand and vermiculite (1:1:1 volume-based). After 56 days of growth for root establishment, salt treatments were applied every 3 days during watering over 28 days to progressively build up soil salinity and to avoid salt shock response (Shavrukov, 2013). The four treatments were: tap water control (EC_water_=0.3 mS cm^-1^), 51mM NaCl (EC_solution_=6.5 dS m^-1^) and 26mM Na_2_SO_4_ (EC_solution_=6.7 dS m^-1^), both considered moderately saline, and 51mM Na_2_SO_4_ (EC_solution_=13.4 dS m^-1^), considered highly saline (Mirck and Zalesny, 2015). The treatments were arranged in a randomised block design with seven replicate plants (seven independent pots, n=7) per treatment (Fig.1). Leaves, stems, and roots were sampled after 28 days of treatment (84 days of growth). A representative subsample of each fraction was immediately flash-frozen in liquid nitrogen and stored at -70°C for metabolomic analysis.

Shoot heights and numbers were measured weekly. Fresh weight (FW) at harvest, oven-dried biomass weight (DW) and moisture content were measured for each organ. The LICOR-600 porometer/fluorometer was used to measure photosynthetic parameters, including stomatal conductance and the quantum efficiency in light of photosystem II ΦPSII (Genty et al., 1989). Soil salinity was inferred at harvest from electroconductivity of the supernatant solution from a soil:water mixture (1:1) and converted to the electroconductivity of saturated soil extract (EC_e_) using the correlation EC_e_=2.23 EC_(1:1)_-0.58 (R^2^=0.98) (Sonmez et al., 2008).

Elemental analysis of roots, stems, and leaves was performed on ground dry biomass (<500 μm) by Natural Resources Analytical Laboratory at University of Alberta (AB, Canada). Briefly, 0.1±0.02 g of sample was digested overnight, with 5 mL of Trace Metal Grade HNO_3_ using the Microwave Accelerated Reaction System (method EPA 3051a (modified)), diluted to 25 mL with milliQ water and analysed for mineral nutrients (Al, B, Ca, Cu, Fe, K, Mg, Mn, Mo, Na, Ni, P, S, Zn) using a Thermo iCAP6300 Duo inductively coupled plasma-optical emission spectrometer (ICP-OES). The anions Cl^-^ and SO_4_^2-^ were determined by extracting 0.2±0.01 g of dry sample with 50 mL of 2% acetic acid using a reciprocal shaker for 30 min, followed by a 0.45 μm PTFE syringe filtration. Chloride and sulphate were measured using the ferrithiocyante colorimetric method (EPA Method 325.2) and the barium chloride turbidimetric method (EPA Method 375.4), respectively.

### 3.2 Extraction of metabolites

Metabolites from willow roots, stems and leaves were extracted according to De Vos et al. (2007) with some modifications. Briefly, plant tissues were freeze-dried for 72 hours and homogenised using a precooled bead mill Tissuelyser II (Qiagen) at 30 Hz, for 2-4 cycles of 60 seconds each. From the dry tissue powder, 50±0.5 mg was suspended in 1 mL of methanol:water (75:25%v/v), also containing the internal quantitative standard resorcinol at 5 mmol L^-1^. Samples were vortexed for 10 s, sonicated in a 135 W/42 Hz ultrasound bath for 10 min at room temperature and centrifuged at 11,000xg for 2 min at 20°C. The supernatant was filtered through a 0.2 μm centrifuge filter, diluted by 4 with methanol 75% and placed in the LC-MS/MS autosampler conditioned at 4°C.

### 3.3 Liquid chromatography-tandem mass spectrometry (LC-MS/MS) acquisition

Chromatographic separation was performed using an Agilent 1260 Infinity system, where 10 μL of extract was injected onto a Zorbax Eclipse Plus C18 column (4.6 × 100 mm, 3.5 μm) at 30 °C. Mobile phase was solvent A (water, 5% methanol, 0.1% formic acid) and solvent B (methanol, 0.1% formic acid) with a flow rate of 0.4 mL min^−1^ and an 80-minute elution gradient: 100% A hold for 20 min, then a linear increase from 0% to 100% B over 50 min and 100% B hold for 10 min. This gradient allowed detection of highly water-soluble (polar) compounds within the first isocratic phase. Untargeted full-scan (100–1,300 Da) MS acquisition was performed on extracts from all three organs (roots, stems and leaves) of each of the seven replicates (84 samples total), blanks and quality control samples using an Agilent Q-TOF 6530B mass spectrometer equipped with an Agilent Jet Stream ion source operating in positive ion mode (ESI (+)). Gas temperature was 300 °C, drying gas flow was 5 L min^-1^, and the nebuliser pressure was 45 psig. Sheath gas temperature was 250 °C with a gas flow of 11 L min^−1^. Data-dependent MS/MS acquisition was performed on three biological organ replicates per treatment and acquired at collision energy of 20 and 35 eV. Precursor isolation used an intensity threshold set to 500, with a maximum of 4 precursors per cycle and 3 spectra per precursor released after 0.3 min.

### 3.4 Untargeted metabolite data processing

Raw LC-MS data were converted to mzML files by using msConvert (Chambers et al., 2012) and processed using MZmine 3.3.0 (Schmid et al., 2023) (Fig.1). Background noise cut-off was 1,000 for MS spectra and 0 for MS/MS spectra, with a minimum 5,000 peak intensity threshold for feature detection. Chromatogram deconvolution used the local minimum search algorithm, isotopes were grouped and features alignment across samples was performed with 20 ppm mass tolerance (80% weighting) and 1 min retention time tolerance (20% weighting). Feature quality control filtering applied isotope patterns (minimum two), minimal occurrence of four across all samples, and a maximal blank:sample average ratio (1:3). Features eluting within <0.2 min retention time and with peak shape correlated by >85% (MZmine metaCorrelate algorithm) were assigned to a feature correlation group and used for curation of adducts, complex formation, and in source fragments.

### 3.5 Metabolites annotation and molecular networking

The feature annotation pipeline used unsupervised and supervised searches across multiple databases integrated with cheminformatic tools (Fig.1). An initial unsupervised annotation step used *in silico* prediction with SIRIUS, combining raw formula and structural elucidation CSI:FingerID (Dührkop et al., 2019), as well as compound class annotation with CANOPUS (Dührkop et al., 2021). In parallel, all features were searched against a set of MS and MS/MS databases, including general databases available in the GNPS environment (including GNPS, MassBank and ReSpect) as well as experiment- and organism-specific databases (PlantCyc (Hawkins et al., 2021), CEU mass mediator (Gil-de-la-Fuente et al., 2019), an in-house library for Salicaceae based on literature as well as authentic standards). Annotation candidate information was standardised for comparison across tools, and the best hit for each feature selected after quality assessment and prioritisation based on integrated confidence/accuracy scores, MS/MS alignment scores, and MS match precision (minimal difference between experimental and predicted compound mass, adduct prediction) to generate a list of putative compounds from combined tools outputs.

In a second supervised step, annotation of significantly differentially abundant compounds was refined using data-driven and knowledge-based curation. Data-driven curation uses feature-based ion identity molecular networking (Z. Zhou et al., 2022) based on cosine and/or Spec2Vec metrics (Huber et al., 2021) of spectral similarity within subnetworks, and manual resolution of ambiguities. Knowledge-driven curation takes advantage of published compound lists and spectra from comparable experiments and plant species not deposited in public databases to improve annotation.

Each putative compound annotation is assigned a confidence level (1-4) (Schymanski et al., 2014; Sumner et al., 2007) and is reported in supporting information – Table S4, following best reporting practices (Alseekh et al., 2021). Chemical taxonomy was assigned to metabolites using the Natural Product Classification system (Kim et al., 2021). Molecular networks were generated in the GNPS analysis environment (Nothias et al., 2020) and visualisation used cosine metrics (at 0.6 threshold) within Cytoscape software (Shannon et al., 2003).

### 3.6 Statistical analysis

Physiological and metabolomic data were processed using R (R Core Team, 2022) and statistical significance was set at a Benjamin-Hochberg (BH) false discovery rate (FDR) adjusted p-value threshold of <0.05. Physiological comparisons between treatments used a two-way ANOVA, followed by a Tukey HSD *post hoc* test (n=7, except n = 3 for mineral element analysis). Metabolite peak areas were log-transformed and normalised using EigenMS (Karpievitch et al., 2014) prior to statistical analysis (Fig.1). To identify differentially abundant (DA) metabolites, pairwise comparisons of peak area for each treatment (n=7) were performed against controls using either a t-test or Wilcoxon rank sum test with FDR (BH) control, depending on whether assumptions of normality and equal variance were met, or through detection as structural zeros (He et al., 2014) (metabolites detected ≥4 replicates of one condition, and not detected in any replicates of the compared condition). For metabolite comparison across organs, a Kruskal-Wallis test was first performed before *post hoc* pairwise comparisons between each organ pair (Wilcoxon rank sum tests with FDR control). The core metabolome was tentatively defined as compounds that did not significantly vary across control organs and meet a metabolic stability coefficient of variation of peak area below 30%. Principal component analysis (PCA) was performed using MixOmics package (Rohart et al., 2017) and PERMANOVA on Bray-Curtis distances was used to test significant differences between groups.

## 4 Results

### 4.1 Physiological effects of salt treatments and mineral elements accumulation

After 28 days of salinity treatments, soil electrical conductivity (EC_e_) was 0.9±0.2 dS m^-1^ in controls (Fig.2a) and was significantly higher in both moderate NaCl and Na_2_SO_4_ treatments, at 6.5±0.2 and 5.5±0.5 dS m^-1^ (Tukey HSD, α<0.05). High Na_2_SO_4_ significantly increased soil conductivity further to 9.1±0.5 dS m^-1^.

Willows maintained a healthy phenotype throughout the experiment (Fig.2b) and salt treatments did not induce significant differences in aboveground growth rates or organ biomass yields compared to controls (Supporting information Fig.S1-S3), with total biomass averaging 131±5 g dry matter (DM) (Fig.2c). Quantum efficiency ΦPSII ranged from 0.44-0.61 and was not significantly different between treatments (Fig.2d). Salt-treated willows showed significant reductions in stomatal conductance to water vapor (g_sw_) ranging from 0.13-0.19 mol m^-2^ s^-1^ (Fig.2e), compared to g_sw_ of 0.29±0.04 mol m^-2^ s^-1^ in control willows. Leaf moisture content (MC) was 1.97±0.07 g_H2O_ g_DM_^-1^ in controls and significantly increased to 2.28±0.10 g_H2O_ g_DM_^-1^ under moderate NaCl treatment. Moderate Na_2_SO_4_ resulted in intermediate leaf MC, at 2.09±0.04 g_H2O_ g_DM_^-1^, while high Na_2_SO_4_ treatment significantly decreased compared to both moderate salt treatments, measuring 1.75±0.04 g_H2O_ g_DM_^-1^ (Fig.2f). None of the treatments induced significant differences in stem moisture content, which was maintained at an average of 1.34±0.03 g_H2O_ g_DM_^-1^.

Total sodium content was 2.78±0.1 mmol(Na) in controls, 9.4±0.3 mmol(Na) in moderate NaCl, 10.1±0.1 in moderate Na_2_SO_4_ and 13.0±0.1 mmol(Na) in high Na_2_SO_4_-treated willows (Fig.3a). Comparison at the organ scale (Fig.3b) revealed the highest sodium concentration in roots, 3-fold higher than controls in all salt treatments. Total chloride content was significantly higher in NaCl treatment, with 26.3±2.1 mmol(Cl) as compared to an average of 6.8±0.2 mmol(Cl) for controls and Na_2_SO_4_ treatments. In NaCl treatment, leaves had the highest concentration of Cl at 575.4±26.5 µmol(Cl) g_DW_^-1^, representing a 3.4-fold increase compared to controls; while accumulation in stems and roots was below 190 µmol(Cl) g_DW_^-1^, but still 4.3 to 5.5-fold higher than controls. Total sulphur content was significantly higher in trees of both Na_2_SO_4_ treatments, at similar 14.1±0.5 and 15.7±1.9 mmol(S), as compared to 7.4±0.4 mmol(S) in controls and 8.5±0.2 mmol(S) in NaCl. Sulphur concentration significantly increased from 84.3±6.1 µmol(S) g_DW_^-1^ in control roots to 147.9±6.1 µmol(S) g_DW_^-1^ in Na_2_SO_4_-treated roots and from 161.0±11.1 µmol(S) g_DW_^-1^ to 229.1±7.5 µmol(S) g_DW_^-1^ in leaves. At the whole plant scale, the proportion of inorganic sulphur did not vary significantly across treatments, averaging 27% of the total sulphur present as inorganic SO_4_^2-^ (Supporting information – Fig.S5). Total potassium content was 21.3±0.6 mmol(K) in controls and significantly higher in both moderate and high Na_2_SO_4_ treatments, with similar content of 28.9±0.2 and 28.4±2.3 mmol(K) respectively. Potassium concentration tended to decrease in salt-treated roots but increased in salt-treated leaves compared to controls. Across all four treatments, there were no significant variations in total phosphorus, calcium and magnesium content (Fig.3a). However, calcium concentration in leaves significantly decreased in both Na_2_SO_4_-treated plants compared to controls while leaf magnesium concentration significantly increased in NaCl-treated plants only, compared to controls (Fig.3b).

### 4.2 Organ partitioning of the Salix metabolome

After 84 days of growth, untargeted metabolomic profiling of control willows, without salt treatment, revealed a total of 5,043 metabolic compounds across all three organs, with 2,063 compounds in roots, 2,950 compounds in stems, and 3,601 compounds in leaves (Fig.4a). Shannon and inverse Simpson indices showed a significant sequential increase in metabolic diversity from roots to stems to leaves (Fig.4b). Principal Component Analysis (PCA) separated samples based on organ (Fig.4c), PC1 and PC2 explaining 52% and 26% of variance, respectively, with significant differences between groups (*PERMANOVA*, p-value < 0.05). Leaves and stems exclusively shared 481 compounds, stems and roots exclusively shared 371 compounds while leaves and roots exclusively shared 83 compounds (Fig.4d).

A total of 1318 compounds were present in all three organs (Fig.4d), representing 73-96% of cumulative peak area in each organ (Fig.4e). Differential abundance analysis between organs showed that, of these shared compounds, 278 did not significantly differ. They were defined as “core metabolites” and included 29 flavonoids, 14 phenolic acids, 12 phenylpropanoids, 22 small peptides, 11 tryptophan alkaloids, 9 pseudoalkaloids, 10 nucleosides, 8 saccharides, 8 monoterpenoids and 4 fatty acids and conjugates (Fig.4f). High-intensity compounds included adenosine, glutamate, 12-tetradecenoic acid, myrciacitrin III, apigenin 7-O-neohesperidoside, kaempferol 3-O-glucoside (astragalin), salicortin, salicin, benzaldehyde, 4-o-caffeoylshikimic acid, secoxyloganin, 7-O-p-coumaroyl loganin, nicotinamide, galactosylglycerol and fructose (Supporting information - Tables S3,S4).

Conversely, 3,259 compounds were organ-specialised metabolites with significantly increased abundance in one organ, including 1,719 compounds exclusively detected in leaves, 780 in stems and 291 in roots. Leaf-specialised compounds were dominated by the terpenoid pathway (Fig.4g), with a high number of (seco)iridoid monoterpenoids, diterpenoids, steroids, (apo)carotenoids, as well as the alkaloids pathway, particularly tyrosine and tryptophan derived alkaloids. Stem-specialised compounds were dominated by phenylpropanoids, lignans and fatty esters whereas root-specialised compounds were dominated by saccharides and amino acids glycosides. Each organ also had a highly specific set of flavonoids and small peptides. Interestingly, 19% of root-specialised compounds were highly water soluble (polar), compared to 5% of stem-specialised and 10% of leaf-specialised compounds (Supporting information – Fig.S6).

### 4.3 Salix metabolomic response to moderate and high Na_2_SO_4_

Principle component analysis (PCA) separated samples by control, moderate and high Na_2_SO_4_ treatments in roots *(PERMANOVA*, p-value <0.05), with PC1 explaining 17% and PC2 explaining 11% of the variance, while no separation was observed between treatments in stems and leaves (Fig.5a). Moderate Na_2_SO_4_ led to 571 differentially abundant (DA) compounds across the whole plant, including 218 in roots, 105 in stems and 248 in leaves (Fig.5b,d). Comparatively, high Na_2_SO_4_ led to 976 DA compounds across the whole plant, including 574 in roots, 131 in stems and 271 in leaves (Fig.5c,d).

Taken together, the Na_2_SO_4_-salt response involved 989 DA compounds distributed across the main pathways as 260 shikimates/phenylpropanoids, 133 alkaloids, 84 carbohydrates, 79 amino acids/peptides, 73 terpenoids, 51 fatty acids and 25 polyketides (Fig.5e). In response to moderate Na_2_SO_4_ treatment, DA compounds in roots and stems were mostly depleted (58-60%), while DA compounds in leaves were mostly enriched (85%) (Fig.5d,h). Higher Na_2_SO_4_ induced a larger metabolite enrichment across most pathways, except for further depletion in carbohydrates, while stems maintained a reduced metabolite pattern and leaves maintained increased metabolites across the major pathways.

Over the whole plant, 327 compounds were consistently altered in both moderate and high Na_2_SO_4_ treatments, representing 21% of DA compounds in roots, 24% in stems and 53% in leaves (Fig.5f). This consistent response was further reflected in a R^2^=0.93 correlation between the fold changes (log_2_) moderate and high Na_2_SO_4_ treatments compared to controls (Fig.5g).

In roots, moderate Na_2_SO_4_ reduced 45 compounds within shikimates/phenylpropanoids pathway, including 15 flavonoids, 9 phenolic acids and 5 lignans (Fig.5h). Comparatively, high Na_2_SO_4_ treatment increased 113 compounds from that pathway, including 52 flavonoids, 12 phenolic acids, 12 phenylpropanoids as well as 10 lignans. These included enrichment of important compounds such as 4-coumaroyl shikimate, 5-hydroxyferulate, procyanidol, catechin 3-O-(1-hydroxy-6-oxo-2-cyclohexene-1-carboxylate), 8-hydroxypinoresinol 4’-glucoside, guaicylglycerol-coniferyl ether, glucose-conjugated salicylic acid, and salicin. Six sphingolipids, including phytosphingosine and 4-hydroxysphinganine, were consistently depleted by both Na_2_SO_4_ concentrations, while 4 and 10 saccharides significantly reduced under moderate and high Na_2_SO_4_, respectively. In contrast, 8 nucleosides and 20 small peptides increased only under high Na_2_SO_4_, as did 8 pseudoalkaloids, 8 tryptophan alkaloids, 6 tyrosine alkaloids, 12 monoterpenoids, 11 steroids and 9 triterpenoids. In stems, moderate and high Na_2_SO_4_ treatments enriched 3 and 7 small peptides, such as aspartate and proline, while 7 and 8 small peptides were depleted, respectively. Additionally, 4 and 14 saccharides were depleted, including glucose and trehalose (Fig.5h), whereas 14 and 13 saccharides were enriched in leaves. Common, dose-responsive compounds in leaves were dominated by 49 shikimates/phenylpropanoids, encompassing enrichment of 9 cinnamic acid derivatives, 5 salicinoids and 4 other simple phenolic acids, 4 flavonols and 4 chalcons. Commonly regulated metabolites also included 25 alkaloids and 20 amino acids/peptides, with the enrichment of 7 pseudoalkaloids (5 purine and 2 pteridine alkaloids), 3 tryptophan alkaloids and 15 small peptides (Fig.5h).

A total of 179 DA compounds altered by one or both concentrations of Na_2_SO_4_ (12% of all Na_2_SO_4_ metabolomic changes) contained sulphur, corresponding to 101 distinct compounds. Roots significantly accumulated 13 and 45 sulphur compounds (S-compounds) under moderate and high Na_2_SO_4_ respectively, but significantly reduced 4 and 20 S-compounds (Fig.5i). Additionally in leaves, 36 S-compounds were increased under one or both Na_2_SO_4_ concentrations, compared to 11 depleted S-compounds. Shared enriched S-compounds included dicarboxylic acid such as 2-(2-sulfoethyl)propanedioic acid or polyamines, such as N’-hydroxy-5-(2-methoxyethyl-sulfamoylamino) pentanimidamide, as well as stilbenes, such as resveratrol 4-sulfate. High Na_2_SO_4_ depleted S-containing compounds ranged from small peptides, such as threonine to larger alkaloids such as C_19_H_16_N_8_O_5_S (CHEMBL164867). Notably, a subgroup of 16 DA S-compounds were in the same structural cluster (Fig.5j).

### 4.4 Salix metabolomic response to NaCl treatment

Comparing metabolomes from controls with NaCl-treated willows, PCA revealed a significant separation between control and NaCl treatment in roots (*PERMANOVA*, p-value <0.05), partial overlap in stems (*PERMANOVA*, p-value < 0.05) and no observed separation in leaves (Fig.6a). NaCl treatment led to DA in 1,063 DA compounds, with 475 in roots, 481 in stems and 107 in leaves (Fig.6b). Among these, 768 DA compounds were unique to one organ, while 135 were shared to two or all organs (Fig.6c). Notably, 98 DA compounds were exclusively shared between roots and stems and 24 were significantly depleted in all three organs due to NaCl treatment.

Across all organs, NaCl-responsive metabolites comprised 213 DA compounds within the shikimates/phenylpropanoids pathway, 118 alkaloids, 88 carbohydrates, 74 amino acids/peptides, 59 terpenoids, 48 fatty acids and 12 polyketides (Fig.6d). While shikimates/phenylpropanoid pathway compounds were the most diverse, carbohydrates represented the highest proportional change within a pathway, constituting >33% of total carbohydrates (Supporting information – Fig.S7). This was driven by significant depletion in 28 carbohydrates across multiple organs, including glucose, fructose, sucrose and valienol 1,7-diphosphate, as well as organ-specific depletion of 36 carbohydrates (Fig.6e,f). Tricarboxylic acid cycle intermediates, such as malate, citrate, 2-oxoglutarate, were also depleted across the entire plant as well as phosphorylated compounds, such as phosphocholine, phosphonoacetaldehyde, 1,7-diphospho-1-epi-valienol and 2-C-methyl-D-erythritol-2,4-cyclodiphosphate (Supporting information – Fig.S8).

Within organs, leaves had a limited response to NaCl treatment (Fig.6b,c), but depleted compounds included 2-oxoglutarate, malate, aspartate and choline, while accumulated compounds included populoside C, (indol-3-yl)acetyl-myo-inositol L-arabinoside (IAA derivative) and spermidine (Supporting information – Tables S3,S7). Stems and roots had a larger metabolic response, markedly within the shikimate/phenylpropanoid pathway, altering 115 and 108 compounds respectively (Fig.6e). This large group of differentially abundant phenolic compounds was primarily associated to flavonoids, phenylpropanoids, lignans and phenolic acids (including salicinoids). While enriched flavonoids number were similar in stems and roots, 20 and 23 DA compounds respectively, stems had more depleted flavonoids than roots, with 23 and 10 DA compounds respectively (Fig.6h). Salicinoids were net enriched in both stems and roots (Fig.6h). Stems had significant enrichment of myricetin 3-O-glucoside, kaempferide 3-O-glucoside, procyanidol, salicin and grandidentatin, alongside the depletion of eriodictyol 7-glucuronide, luteolin 7-(6’’-benzoylglucoside, ramontoside and a hydroxycyclohexenoyl ester catechin derivative. In roots, populin, tremuloidin and benzoyl-β-D-glucoside were enriched, while (iso-)salicin, salicylic acid glucoside, grandidentatin, catechin, procyandin C, pleurostimin 7-glucoside were depleted. Additionally, 21 lignans and phenylpropanoids significantly accumulated in roots, contrasting stems, where 14 lignans and phenylpropanoids were depleted (Fig.6h).

A balanced pattern of metabolomic change was observed in nitrogen metabolism in response to NaCl treatment, with enrichment of 147 DA compounds and depletion of 133 DA compounds. This was particularly evident as 31 small peptides increased across all organs, including proline, tryptophan and glutathione, while 34 small peptides decreased, including acetylproline, threonine and arogenate (Fig.6g). Other increasing N-containing compounds included purine nucleosides like adenosine, alkaloids such as dihydrosanguinarine and kinetin-7-N-glucoside, as well as sphingolipids, such as phytosphingosine and sphinganine. Concurrently, terpenoids were enriched, especially monoterpenoids, with 15 enriched monoterpenoids in stems and 13 in roots (Fig.6i,j).

### 4.5 Salix anion-specific metabolomics response to NaCl and Na_2_SO_4_ (at the same EC)

Comparing the 1,063 and 571 DA compounds altered by moderate NaCl and moderate Na_2_SO_4_ revealed 103 DA compounds commonly regulated in roots, 53 in stems and 7 in leaves (Fig.7a,b). These generalised salinity responsive metabolites included 23 shikimates/phenylpropanoids, 10 alkaloids, 10 carbohydrates, 9 terpenoids, 9 amino acids and peptides and 5 fatty acids (11 other and 70 unknown) (Fig.7b). Commonly enriched compounds included (indol-3-yl)acetyl-myo-inositol L-arabinoside and 2-phenylacetamide in leaves, proline, grandidentatin and fraxiresinol 1-O-glucoside (a lignan) in stems, 2-thioethylidenebis(phosphonic acid), delphinidin 3-O-(6’’-O-malonyl)-beta-D-glucoside-3’,5’-di-O-beta-D-glucosideand and syringin in roots. Commonly depleted compounds included glucose, trehalose in stems, D-altro-heptulose, glycerophophoglycerol, benzoyl-beta-D-glucoside and N-(4-(1-cyanocyclopentyl)phenyl)-2-((pyridin-4-ylmethyl)amino)nicotinamide in roots, as well as d-glycero-l-galacto-octulose in both stems and roots.

Anion specific responses included 764 DA compounds unique to moderate NaCl treatment and 391 DA compounds unique to the moderate Na_2_SO_4_. Both salt treatments induced the accumulation of salt-specific monoterpenoids, as well as phenolic compounds, including flavonoids, phenylpropanoids, and phenolic acids. Moreover, NaCl treatment led to a net decrease of saccharides and nucleosides, whereas Na_2_SO_4_ treatment resulted in their net accumulation (Fig.7c). An additional contrasting effect was observed with sphingolipids, with increased in NaCl but decreased Na_2_SO_4_ treated trees.

An anion-specific effect could also be observed when comparing change in constitutively produced compounds that were core (present and relatively stable between organs) and organ-specialised (significantly different between organs), but also in salt-induced compounds (not detected in controls). In both moderate salt treatments, roots primarily reduced root-specialised compounds rather than core compounds, while significantly enriched compounds were predominantly newly synthesised (salt-induced), rather than root-specialised or core compounds (Fig.7d). Similarly, in stems, reduced compounds were largely stem-specialised rather than core compounds under both salt types, however enriched compounds were predominantly stem-specialised under NaCl, comprising 53 enriched DA compounds as compared to 20 salt-induced and 23 core enriched DA compounds. Conversely, Na_2_SO_4_ treatment resulted in 2 stem-specialised, 5 salt-induced and 3 core enriched DA compounds. In leaves, NaCl treatment reduced 13 leaf-specialised and 4 core DA compounds while enriching 26 leaf-specialised, 20 salt-induced and 4 core DA compounds. In contrast, leaves treated with Na_2_SO_4_ decreased 22 leaf-specialised and 4 core compounds while increasing 5 leaf-specialised and 63 core compounds.

## 5 Discussion

### 5.1 Salix salinity tolerance response

Soil salinity levels are commonly categorised based on EC_e_ as slightly saline (2-4 dS m^-1^), moderately saline (4-8 dS m^-1^), and highly saline (>8 dS m^-1^) (Corwin and Scudiero, 2019; FAO, 1988). Here, the comparison of non-saline control, moderately saline NaCl and Na_2_SO_4_, and highly saline Na_2_SO_4_ (Fig.2a) enabled capture of the anion-specific effects (Cl^-^ and SO_4_^2-^) and severity effects (moderate vs high Na_2_SO_4_) on willow (*Salix miyabeana* ‘SX64’).

**Figure 2:**
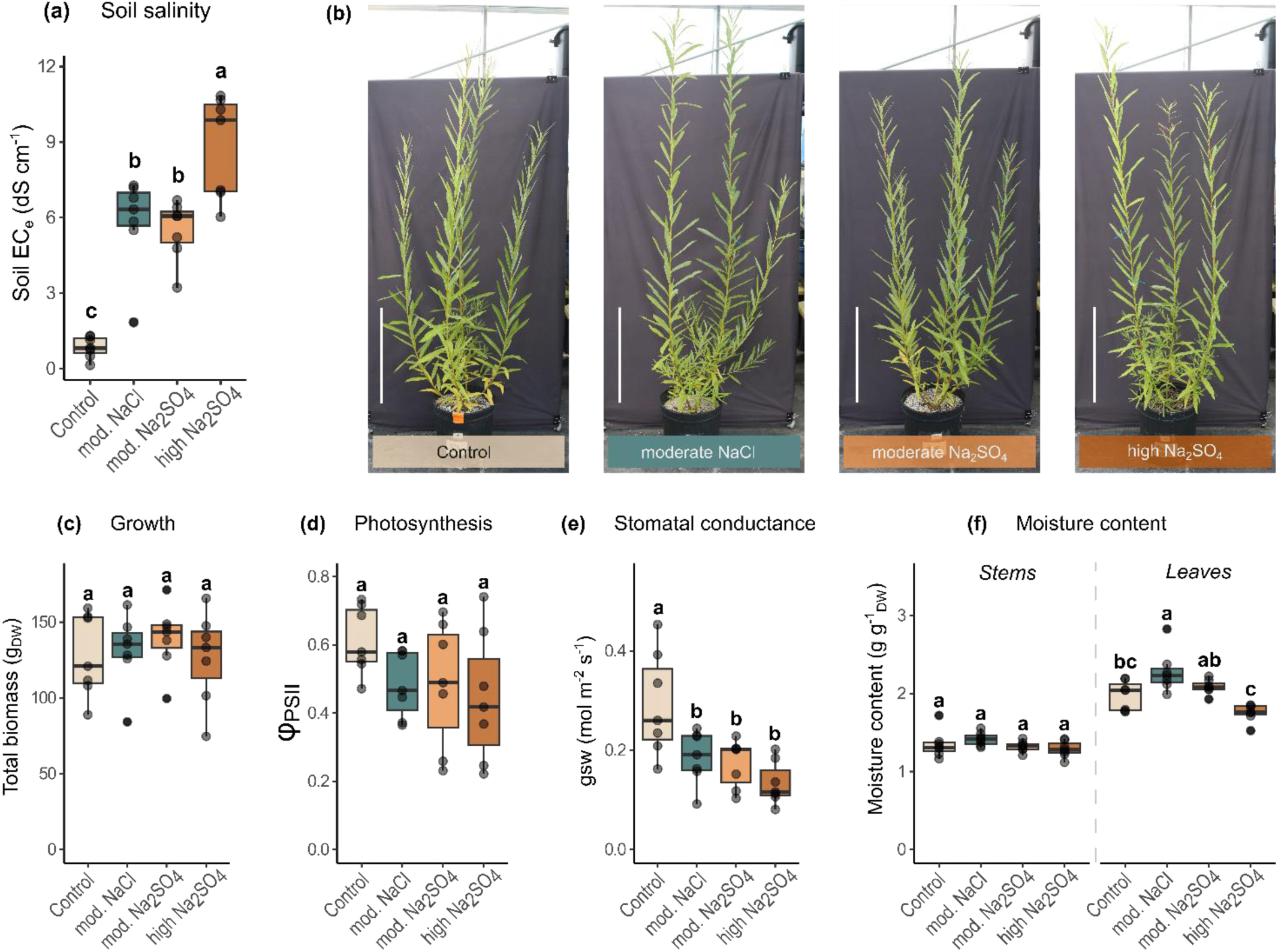
Physiological assessment of salt stress on Salix miyabeana ‘SX64’. (a) Following 56 days of root establishment, salinity in planted pots was progressively built-up over 28 days to reach the final saturated soil electroconductivity (EC), as inferred from a (1:1) soil-to-water extract. **(b)** Representative willow photographs, on harvest day, in response to salt treatments (white scale bars: 50 cm). The impact of moderate NaCl, moderate Na_2_SO_4_ and high Na_2_SO_4_ were quantified on **(c)** total biomass (dry weight), **(d)** photosystem II efficiency evaluated using the LI-COR 600 parameter phiPSII, **(e)** stomatal conductance gsw, **(f)** stems and leaves moisture contents. Boxes represent median and quartiles while whiskers extend to the largest value within 1.5 times the inter-quartile range (IQR) from the hinge. Dots represent individual plants (n=7). Significant differences (p<0.05) indicated by letters were determined by two-way ANOVA followed by Tukey’s post hoc test.

Photosynthetic capacity and biomass productivity were maintained compared to water controls, even at high soil salinity levels of 9 dS m^-1^ (Fig.2c,d), supporting the suitability of willow for phytoremediation. This aligns with research by Hangs et al. (2011), reporting salinity tolerance in 37 willow varieties, where most cultivars could tolerate moderate soil salinity and the most salt-tolerant cultivars maintained yield under high soil salinity. Leaf moisture content increased under moderate salinity, alongside a >35% reduction of stomatal conductance (Fig.2e). This indicates reduced transpiration accompanied by osmotic adjustment to sustain turgor, potentially mitigating ion toxicity through absorbed salt dilution (Munns et al., 2020b; Nguyen et al., 2017). However, subsequent lower leaf moisture content suggests this water balance might not be maintained as salinity severity increase (Fig.2d).

Salt treatments did not reduce the acquisition of essential nutrients and even increased potassium uptake alongside salt ions, although these ions were selectively partitioned (Fig.3). Sodium accumulation was predominantly restricted to the roots and influx did not increase linearly with external concentration, suggesting a robust sodium exclusion mechanism (Fig.3a). Such exclusion can occur through root cell wall Na^+^ binding (Byrt et al., 2018) or competing uptake of K^+^ (Hauser and Horie, 2010), as proposed in the salt tolerant *Salix interior* (Major et al., 2017) and *Salix eriocephala* ‘LAR’ (Huang et al., 2023). In contrast, the high chloride uptake and transport throughout the plant (Fig.3) can be facilitated by passive anion channels (Skerrett and Tyerman, 1994), resulting in leaf chloride concentrations here reaching the upper optimal level for non-halophytes (∼30-560 μmol(Cl) g_DW_^-1^ (Geilfus, 2018)). This accumulation, coupled with increased leaf water content, suggests that Cl^-^ may serve as a readily available osmolyte, potentially conferring osmotic benefits similar to those observed in *Nicotiana tabacum* (Franco-Navarro et al., 2016). However, given that cytosolic chloride toxicity is thought to occur at concentrations of 5-20 mmol(Cl) L(cytosol)⁻¹ (Teakle and Tyerman, 2010), the accumulated Cl^-^ may also require compartmentalisation into vacuoles, apoplasts, or older leaves to mitigate cellular damage and optimise metabolic burden. In comparison to chloride, sulphate uptake was more constrained but still resulted in increased transport throughout the plant, where it was either stored as SO_4_^2-^ or metabolised into organic sulphur compounds.

**Figure 3:**
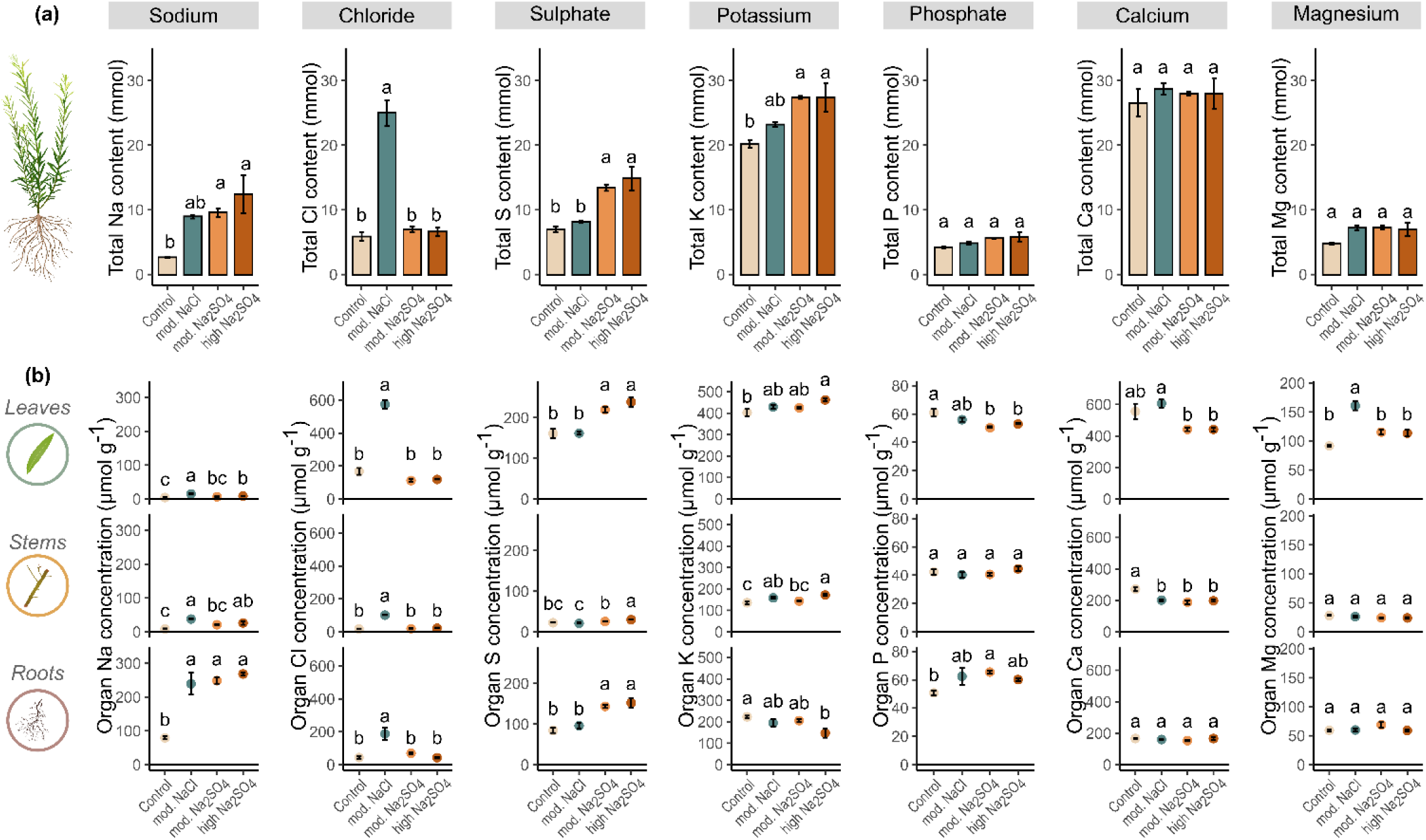
Elemental analysis of mineral nutrients. Elemental analyses include sodium (Na), chloride (Cl), sulphur (S), potassium (K), phosphorus (P), calcium (Ca), and magnesium (Mg). **(a)** Total element content (mmol) combined from the leaves, stems and roots presented. The bars indicate mean values within each organ ± standard error (n=3). **(b)** Element concentration within a single organ (μmol g_DW_^-1^) is shown. The dots represent mean values ± standard error (n=3). Letters indicate significant differences (p<0.05) between control, moderate NaCl, moderate Na_2_SO_4_, and high Na_2_SO_4_ treatments and were determined using two-way ANOVA followed by Tukey’s post hoc test. Mineral elements ratios (K^+^/Na^+^; Ca^2+^/Na^+^; K^+^/Ca^2+^) are presented in Supporting information – Fig.S4.

The maintenance of biomass yields despite reduced stomatal conductance, which can lead to decreased carbon assimilation (Munns et al., 2020a), suggests effective metabolic plasticity and ion management strategies, making this experimental system suitable for assessing willow metabolomic tolerance mechanisms without the confounding metabolic effects associated with severe salt-induced physiological damage.

### 5.2 High organ-specific diversity in the Salix metabolome

Untargeted metabolomic analysis of control trees revealed substantial diversity across the three main vegetative organs (roots, stems and leaves), with over 5,000 compounds detected, approximately 62% of which could be annotated at confidence levels 1-3 (Sumner et al., 2007), providing a detailed overview of the willow metabolome at whole plant scale (Fig.4). Previous metabolomic studies in single organs of *Salix* have captured up to 1,639 metabolites in bark (J. Zhou et al., 2022), 2,735 (Kaling et al., 2015), 600 (Jia et al., 2020) and 2,440 (Aliferis et al., 2015) metabolites in leaves, and 800 metabolites in roots (Xia et al., 2021). By integrating analyses of multiple plant organs, a group of common metabolites across organs were identified, representing 26% of metabolomic diversity and over 70% of the putative abundance (estimated from peak area) (Fig.4d). Of these, 287 metabolites had stable abundance across organs and represent its potential plant-wide basal, or “core”, metabolism (Fig.4f); a recent metabolomic concept that can also denote evolutionarily conserved metabolites across species (Drapal et al., 2022; Dussarrat et al., 2022; McLaughlin et al., 2023). Many of these core metabolites were associated with central pathways like glycolysis, TCA cycle derivatives, and amino acid metabolism, reflecting primary metabolism across plant tissues. Others included bioactive phenolics found at high levels in Salicaceae, such as characteristic salicylates (e.g. salicin, salicortin) associated to antioxidant, antimicrobial and anti-herbivory functions in willow (Boeckler et al., 2011; Brereton et al., 2017; Meier et al., 1988; Pobłocka-Olech et al., 2007; Tyśkiewicz et al., 2019; Volf et al., 2015).

**Figure 4:**
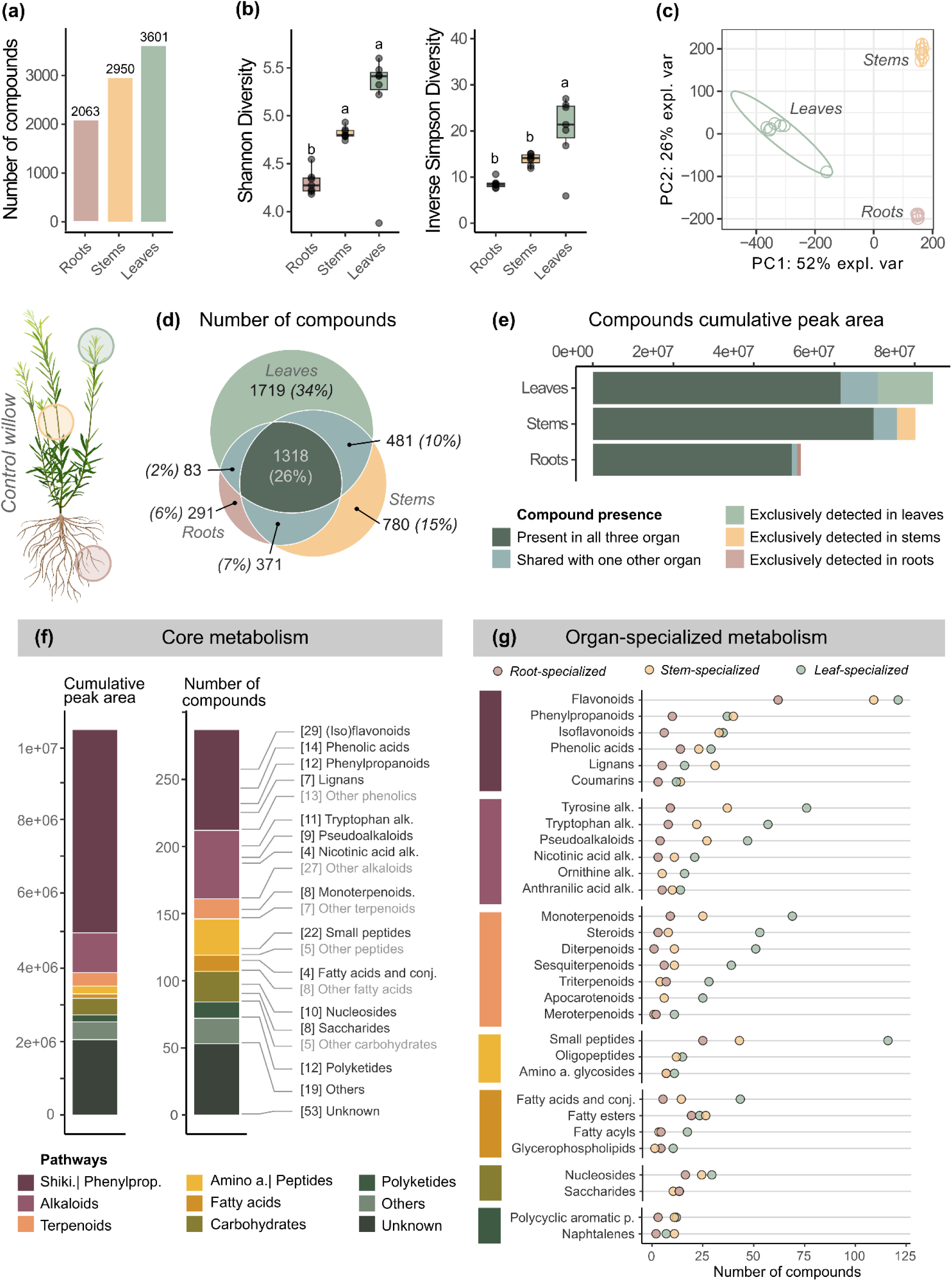
Willow metabolic composition. Organs (leaves, stems and roots) were separately collected from control willows and analysed for metabolomic profiling. **(a)** Total number of consistently detected metabolites within each major organ (present in at least 4 replicates out of 7). **(b)** Comparison of organ metabolic richness using Shannon and Inverse Simpson diversity indices (n=7), different letters indicate significant differences (ANOVA, Tukey HSD, p<0.05). **(c)** Principal component analysis depicting dissimilarity in organ composition based on Bray-Curtis distances, ellipses are for standard deviation. **(d)** Venn diagram illustrating the distribution of metabolites across organs as unique to one or shared between multiple. **(e)** Cumulative peak areas of metabolites present in each organ, categorised based on their exclusive presence in the respective organ or shared across multiple organs. **(f)** Cumulative peak area and number of core metabolites (compounds present across the entire plant, with no significant variation of abundance between organs) classified per metabolic pathways, with composition details of main superclasses. **(g)** Count of compounds significantly higher in one organ compared to the other two (ANOVA, Tukey HSD, p<0.05), revealing dominant organ-specialisation of some metabolic superclasses.

Beyond these common metabolites, willow organs had clear metabolic differences (Fig.4g). Leaves contained the most diverse metabolome, aligning with leaf function as a source of complex photosynthates. The leaf metabolome included diverse terpenoids involved in light harvesting, development, defence, and signalling (Pichersky and Raguso, 2018). Stems accumulated lignans and phenylpropanoids, reflecting woody tissue functions of structure and storage (Boeckler et al., 2011; Köhler et al., 2020). Additionally, the metabolome of stems had a large overlap with both leaves and roots, underscoring their function as conduits for biochemical transport and dialog between aerial and belowground processes (Fig.4d). Comparatively, despite critical functions of water and nutrient absorption, exudation, anchoring structure and storage, roots had the lowest metabolic diversity (Fig.4a,b). Roots were enriched in saccharides and flavonoids, thought to mediate soil interactions like microbial symbiosis (Endo et al., 2021), metal tolerance (Osawa et al., 2011), nutrient uptake (Cesco et al., 2012) and root architecture (Buer et al., 2010). The root metabolome was also distinctly richer in polar metabolites (Supporting information – Fig.S6) which can more readily diffuse through the soil environment within the water vector and could therefore play a role as exudates involved in shaping the rhizospheric biotic (Sasse et al., 2018) and abiotic (Badri and Vivanco, 2009) environment.

These distinct metabolomes reflect the different physiological roles of each organ, describing different metabolic activity and degree of functional specialisation (Li et al., 2016) as well as serving as a baseline to enable understanding of key responses to stimuli such as salt stress.

### 5.3 Sulphate salt induces dose-dependent metabolomic responses in Salix

Sulphate salinity induced extensive organ-specific metabolic reprogramming (Fig.5a-d,h), affecting 10% and 16% of all metabolites at moderate and high concentrations, respectively. Doubling the Na_2_SO_4_ concentration in soil resulted in a twofold increase of metabolic changes in roots, but more limited responses in stems and leaves. This localisation of metabolic response to the roots, which serve as the primary defence barrier against excess ions and osmotic stress, is likely decisive in mitigating salinity stress impact aboveground. Notably, consistent levels of salt ion uptake and accumulation despite increased soil salinity indicate effective ion management (Fig.3), allowing leaves to maintain similar adaptative metabolomic strategies under both sulphate salt treatments (Fig.5f,g). Previous studies reported sulphate salt tolerance in willows (Huang et al., 2020) associated with ion imbalance restriction to roots, similarly preventing physiological impact on leaves at moderate salinity levels.

**Figure 5:**
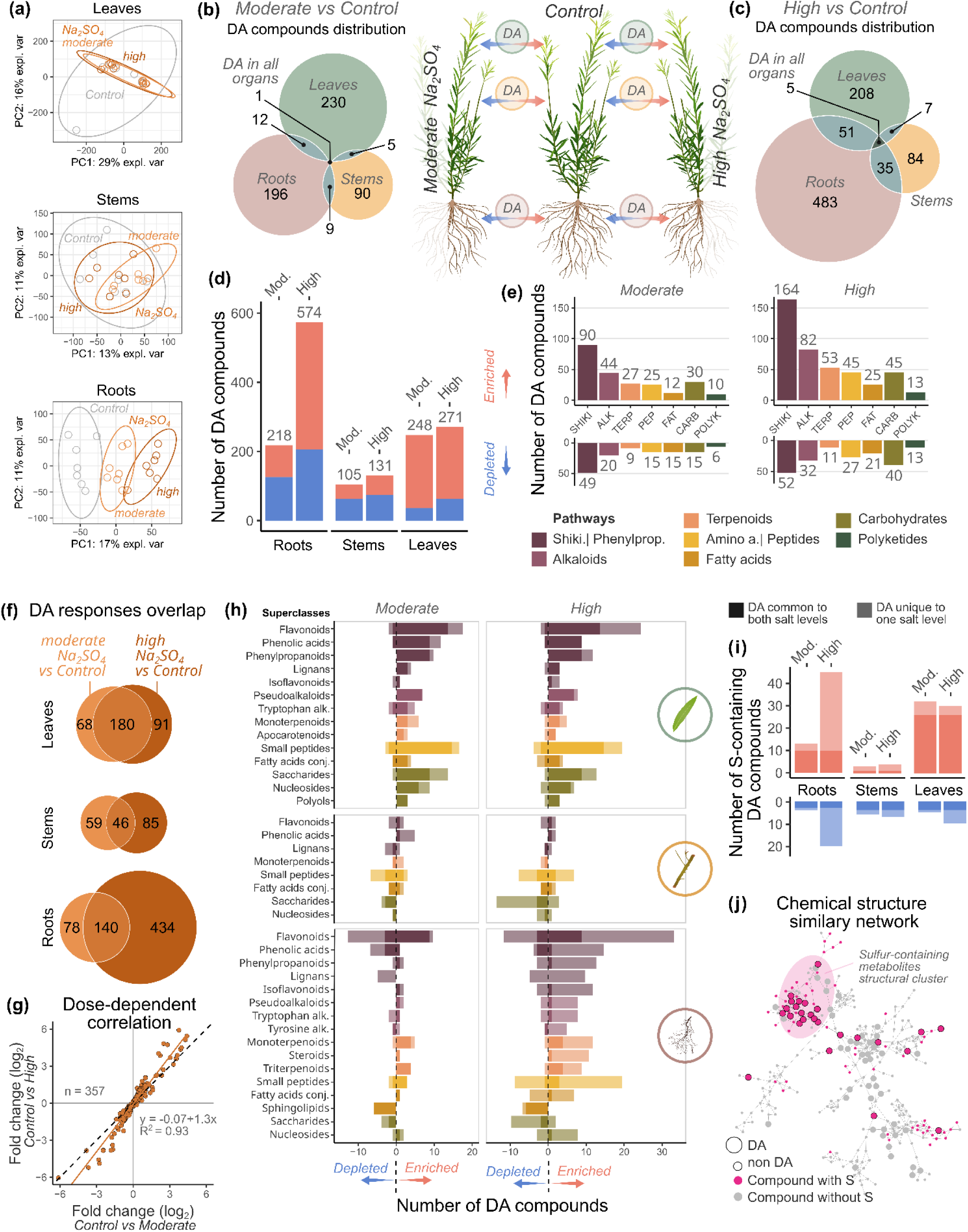
Willow metabolic response to sodium sulphate (Na_2_SO_4_) salt and its variation with dose. **(a)** Principal component analysis (PCA) based on Bray-Curtis distance, illustrating metabolic profile dissimilarity between control, moderate Na_2_SO_4_ and high Na_2_SO_4_ at organ scale (n=7), including standard deviation ellipse (95% confidence interval). **(b)** Organ distribution of differentially abundant (DA) compounds between control and moderate concentration Na_2_SO_4_-treated willows (p<0.05). **(c)** Organ distribution of differentially abundant compounds between control and high concentration Na_2_SO_4_-treated willows (p<0.05). **(d)** Overview of significantly enriched and depleted compounds across organs, comparing DA of control against moderate Na_2_SO_4_ and control against high Na_2_SO_4_. **(e)** Distribution of enriched and depleted DA compounds across metabolic pathways for moderate and high Na_2_SO_4_ treatments compared to controls. **(f)** Metabolic response overlap across both Na_2_SO_4_ concentrations, representing the number of DA compounds common and unique to a single dose DA, at organ scale. **(g)** Dose-dependent correlation of compounds consistently DA across Na_2_SO_4_ levels within the same organ (excluding salt-induced/salt-suppressed compounds with infinite fold change), representing the fold change (log2) of average peak areas (n=7) under moderate Na_2_SO_4_ relative to control against high Na_2_SO_4_ relative to control. **(h)** Sub-selection of main superclasses altered by moderate or high concentration of Na_2_SO_4_ compared to controls, illustrating enriched and depleted compounds distribution at organ scale. DA compounds shared between both treatment levels are highlighted by the darker section of the bars. **(i)** Distribution of enriched and depleted sulphur-containing DA compounds at organ scale, comparing both levels of Na_2_SO_4_. Shared compounds are highlighted by the darker section of the bars. **(j)** Molecular network of compounds MS/MS spectra similarities (cosine >0.6) computed in the GNPS environment and visualised in Cytoscape. Dots are color-coded based on the presence of sulphur (pink represents S-containing compounds) and dot size highlights significantly different compounds (larger dots are DA compounds). A major cluster containing 16 S-containing and structurally similar DA compounds is circled by an ellipse.

**Figure 6:**
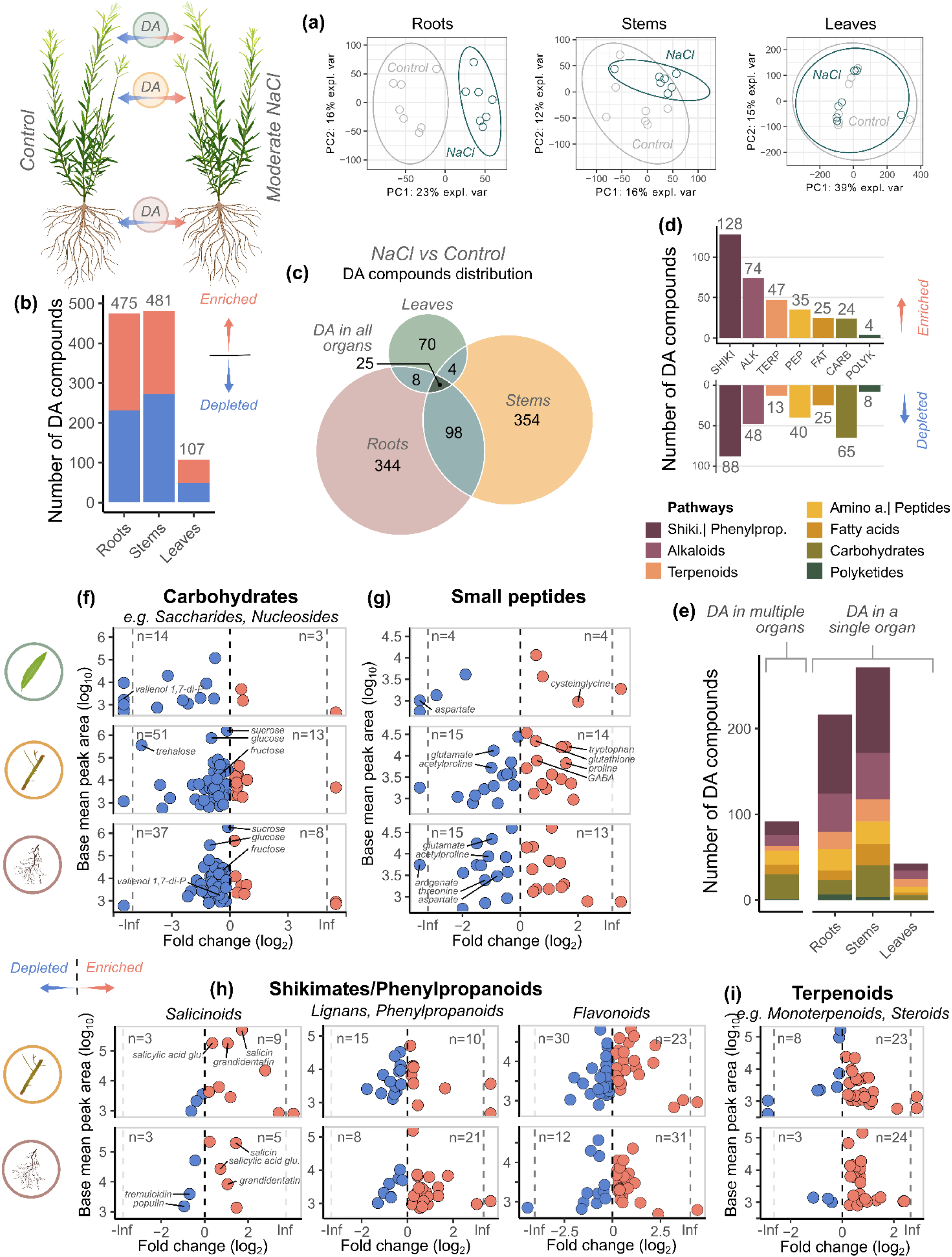
Willow metabolic response to sodium chloride (NaCl) salt. **(a)** Principal component analysis (PCA) based on Bray-Curtis distance, illustrating metabolic profile dissimilarity between control and moderate NaCl at organ scale (n=7), including standard deviation ellipse (95% confidence interval) **(b)** Overview of significantly enriched and depleted compounds across organs. **(c)** Organ distribution of differentially abundant (DA) compounds between control and moderate concentration NaCl-treated willows (p<0.05). **(d)** Distribution of enriched and depleted DA compounds across metabolic pathways. **(e)** Metabolic profile at pathway level of DA compounds shared between multiple organs and unique to roots, stems or leaves. Sub-selection of main pathways or superclasses altered by moderate NaCl compared to controls, illustrating enriched and depleted compounds distribution at organ scale for **(f)** carbohydrates, including saccharides and nucleosides, **(g)** small peptides, **(h)** and 3 major superclass of shikimates and phenylpropanoids pathway: salicinoids, lignans, phenylpropanoids, flavonoids, as well as **(i)** terpenoids, including monoterpenoids and steroids. Dots are color-coded by regulation (blue: depleted, red: enriched by salt treatment compared to control). Base mean area reflects average peak area across all compared samples.

**Figure 7:**
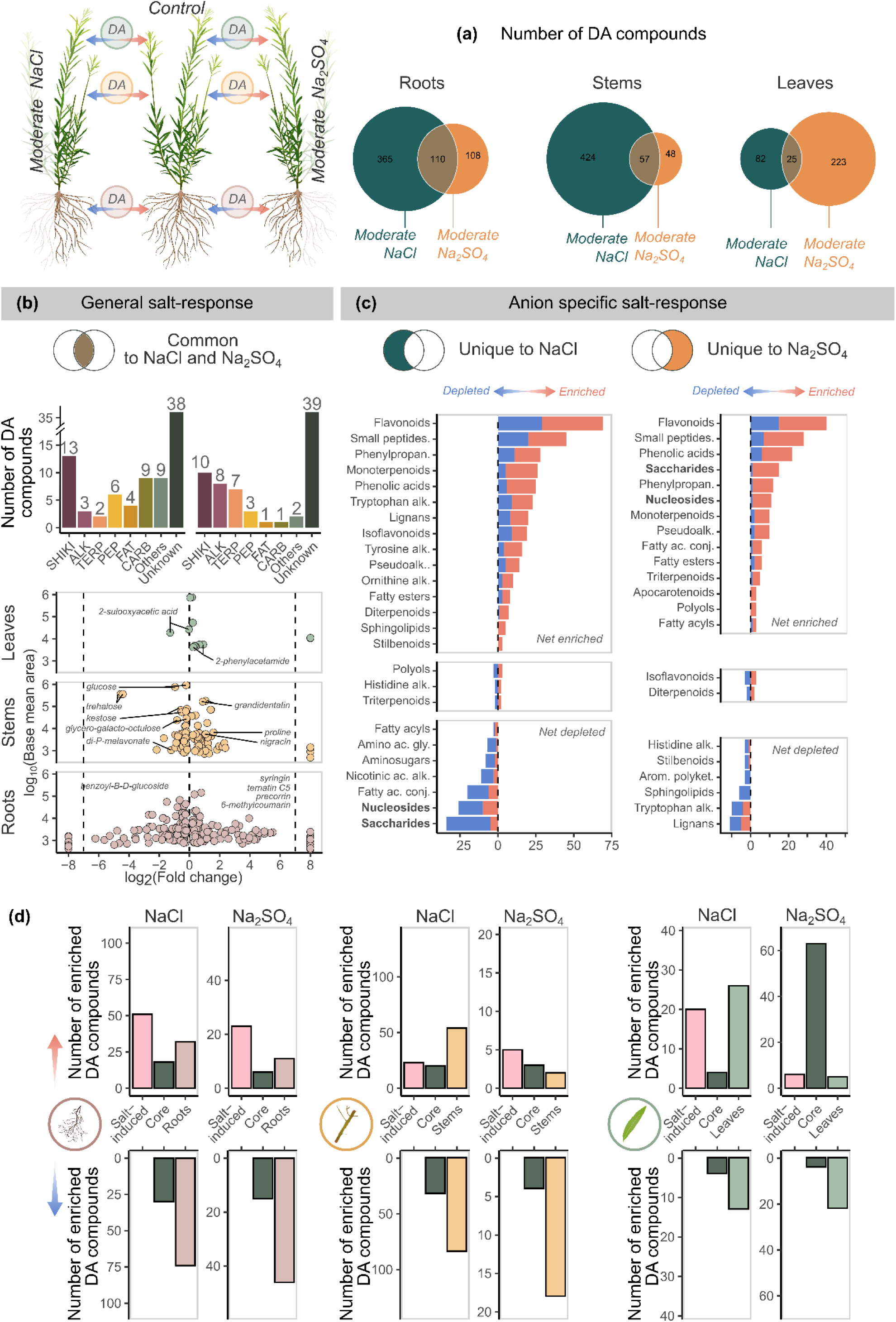
General willow salt response and anion specific tailored responses. **(a)** Venn diagrams showing DA metabolic responses overlap in roots, stems, and leaves under moderate NaCl and Na_2_SO_4_ stress. **(b**) Pathway profile of commonly enriched and depleted compounds in response to moderate salt stress. Fold change graph illustrated enriched and depleted compounds distribution at organ scale. **(c)** Superclass profiles of NaCl- and Na_2_SO_4_-specific responses. Superclasses are categorized as net enriched, equal, or net depleted based on the relative number of enriched versus depleted compounds. Differential patterns for nucleosides and saccharides are highlighted in bold. **(d)** Metabolic response to moderate NaCl and moderate Na_2_SO_4_ categorised based on compounds distribution across the whole plant: salt-induced compounds (not detected in controls), core compounds (present in all organs at stable state in controls), organ-specialised compounds: roots, stems or leaves (present in a higher abundance in a specific organ, based on controls). Bar graphs are split as depleted and enriched compounds to highlight patterning of the DA compounds.

In roots, Na_2_SO_4_ dose differentially effected shikimates/phenylpropanoids, with a reduction of flavonoids, phenolic acids and lignans under moderate level, contrasting their net increase under higher sulphate salinity (Fig.5h). This may indicate a developmental slowdown under mild stress, superseded by a required metabolic burst under more severe salinity levels. The lower osmotic potential associated with higher salinity likely drove the accumulation of lignans (Fig.5h) to enhance structural protection (as lignin precursors) and cavitation protection (Reyt et al. 2021), while accumulation of antioxidative flavonoids can enhance secondary oxidative stress tolerance (Nakabayashi et al., 2014). Under high Na_2_SO_4_, the sugar donor UDP-D-galactose significantly increased alongside phenolics, indicating a disruption in glycosylation and flavonoid bioactivity regulation (Xiao et al., 2014), while increases in iridoid monoterpenoids, such as lamiide, can prevent lipid peroxidation to protect cell membrane integrity (Delaporte et al., 2002). The high diversity of salt-responsive metabolites captured here implies additional functions beyond ROS scavenging, likely contributing to the observed Na_2_SO_4_ stress tolerance. Moreover, the reduction in sphingolipids, which regulate ions through early sensing and signalling, could limit salt uptake below toxic thresholds (Jiang et al., 2019; Liu et al., 2022), coherent with depletion of the essential precursor octanoyl-CoA (Neess et al., 2015; Sonar et al., 2020).

Primary metabolites declined in stems under both sulphur salt concentrations, including reductions in small peptides, saccharides, and nucleosides (Fig.5h). A slowdown of developmental processes and primary metabolism would be expected under salt stress, although there was no accompanying (short-term) impairment to total biomass production (Fig.2c). The comparatively limited disruption of secondary metabolites in stems indicates maintenance of homeostasis and primary resource allocation to support root functions, effectively restricting salt stress propagation.

In leaves, the salt-responsive metabolites were broadly shared between both Na_2_SO_4_ levels, characterised by dose-dependent accumulation, including saccharides, small peptides and precursors of phenolic compounds (Fig.5f,g). Again, illustrating the buffering role of roots in protecting essential carbon fixation functions and energy-related compound production, even under severe salt exposure.

An increase of sulphur-containing compound synthesis under SO_4_^2-^ treatment and upregulation of sulphur assimilation and transport pathways is a characterised plant adaptation mechanism to effectively utilise excess SO_4_^2-^ (Buchner et al., 2004; Davidian and Kopriva, 2010). Here, a substantial enrichment of small peptides occurred in roots under sulphur salinity (Fig.5h), with around half incorporating sulphur. This accumulation of sulphur-containing compounds was consistent throughout the plant at both concentrations (Fig.5i) and was predominantly cysteine and glutathione-based small peptides and some fatty acyl thioesters and alkaloids. The integration of sulphate ions into organic sulphur-containing compounds has the potential to reduce ion imbalance *in planta* during salt stress and could represent a tolerance mechanism in willow. Moreover, the increase of structurally related compounds here (Fig.5i), although their functional information is limited, suggests activation of a bespoke, yet-to-be-described, metabolic response to Na_2_SO_4_. Watanabe et al. (2019) highlighted a need for integrated sulphur-omics to overcome this shortfall in functional understanding of sulphur-containing molecules in plants, of particular relevance to salinity tolerant non-model organisms such as willow.

### 5.4 Salt anions shape Salix metabolomic tolerance responses

The salt response across all treatments involved an extensive proportion of the metabolome (28% of metabolites), reflecting the highly plastic nature of plant metabolism under environmental challenge. While activation of the calcium sensing signalling pathway by Na^+^ (Munns and Tester, 2008) and disruption of the Na^+^/K^+^ ratio (Zhang et al., 2018) are commonly considered to trigger salt responses in plants, anion driven metabolomic responses have been less well explored. Comparison of metabolomic responses to moderate NaCl and Na_2_SO_4_ treatments here, at equivalent soil electroconductivity and sodium concentrations, revealed only 3% of metabolites were consistently responsive, highlighting extensive anion-specific changes (Fig.7).

The common, or generalised, salt metabolite changes were asymmetrically distributed across organs, with ∼15-fold more generalised salt responses in roots than in leaves (Fig.7b). These represent common metabolic strategies to tolerate (anion agnostic) osmotic stress in roots at the root-soil interface and restrict sodium translocation, while leaves might be more susceptible to the (anion dependent) ion toxicity of translocated Cl^-^ and SO_4_^2-^. The generalised salt response included common accumulation of the phenylpropanoid precursor 2-phenylacetamide in leaves (a salt biomarker in the euhalophyte *Suaeda salsa* (Bao et al., 2023)), protective molecules in stems such as the anthocyanin ternatin C5 (Kovinich et al., 2015) and a highly antioxidative coumaric acid conjugate of salicin, grandidentatin (Si et al., 2009), as well as potential cell wall structural components in roots, such as the monolignol precursors syringin.

NaCl treatment, unlike Na_2_SO_4_, induced a comparatively small metabolomic response in leaves (Fig.6a-c). Of the limited responses, a significant accumulation of spermidine, a polyamine known for its acid- and ROS-neutralising properties, could help protect cell membranes, reduce chlorophyll loss and leaf senescence (Gill and Tuteja, 2010b). The auxin conjugate, (indol-3-ylacetyl)-myo-inositol L-arabinoside, also increased, suggesting oxidative stress specific developmental regulation (Bajguz and Piotrowska, 2009; Ludwig-Müller, 2011). A larger change under NaCl salt exposure was the reconfiguration of carbon resources through the shikimate/phenylpropanoids pathway (Fig.6d,h). Specifically, lignans and phenylpropanoids had organ-specific responses: substantial accumulation in roots and significant depletion in stems (Fig.6h). As major lignin precursors, these lignan and phenylpropanoid dynamics could represent the expected structural adaptations within the roots and stems. In particular, lignin deposition can act as a barrier regulating ion uptake and translocation to aerial parts (Byrt et al., 2018; Neves et al., 2010; Reyt et al., 2021).

Flavonoids, whose antioxidative capacity can facilitate localised neutralisation of ROS (Nakabayashi et al., 2014), also had marked organ-specific responses, with substantial accumulation in roots and variable regulation in the stems (Fig.6h). As flavonoids, phenolic acids, lignans and phenylpropanoids all derive from the same substrates (phenylalanine and the pivotal intermediate cinnamic acid), the organ-specific shift suggests a coordinated metabolic response promoting defence in the directly exposed roots while simultaneously restricting the availability and energy expense of these compounds in the stems. By comparison, moderate Na_2_SO_4_ did not induce similar extensive accumulation of phenolic compounds in roots or stems but this response was triggered under higher Na_2_SO_4_ concentration.

Accumulation of osmoprotectants under salt stress is one of the most prevalent and well-described adaptive strategies to reduce cell osmotic potential and stabilise cellular components (D’Amelia et al., 2018; Yang and Guo, 2018). Surprisingly, except for stem proline enrichment under both NaCl and Na_2_SO_4_, the levels of expected organic osmoprotectants, namely hydrophilic and low molecular weight sugars, polyamines or amino acids, were largely reduced in roots and stems under NaCl while accumulating in leaves under Na_2_SO_4_, more distal to the primary site of osmotic stress (Fig.5h, Fig.6f,g and Fig.7c). This response contrasts salt-tolerant poplars which can accumulate soluble sugars as a tolerance mechanism (Jouve et al., 2004; Watanabe et al., 2000) and might be a distinct strategy used by *S. miyabeana* to mitigate osmotic stress through the uptake of inorganic ions, including root restricted Na^+^ and anions (Cl^-^ and SO_4_^2-^) translocation to leaves. Similarly, Ottow et al. (2005) reported preferential sodium uptake by *Populus euphratica*, where Na^+^ served as a readily-available osmoprotectant stored in the apoplast or bound in the cell wall, alongside reduced calcium and soluble carbohydrate levels.

The anion-dependent regulation pattern of osmoprotectants also reflected broader differing metabolic adjustments within central metabolism. Although reduction of stomatal conductance was similar between both salts and photosystem II efficiency (Φ_PSII_) preserved (Fig.2), the impact on foliar glucose and energy-related compounds was distinct. NaCl induced the plant-wide reduction of glucose and multiple TCA cycle intermediates (Supporting information – Fig.S8), representing impaired photosynthetic carbon fixation and decreased flux through glycolysis and the TCA cycle, limiting mitochondrial respiration and overall energy supply. A similar reconfiguration of resources and metabolic processes has been observed under salt stress in barley roots (Wu et al., 2013), where glycolysis was inhibited, and in *Arabidopsis thalia* (Hartmann et al., 2015), where a shift to non-oxidative fermentative energy metabolism can occur. This was coupled with reduced levels of small peptides, nucleosides and other simple carbohydrates like fructose and sucrose in stems and roots (Fig.6f,g and Fig.7c), possibly driven by increased carbon demand for secondary metabolism involved in salt stress tolerance. By comparison, Na_2_SO_4_ induced an increase of glucose in leaves (Supporting information – Fig.S8), alongside an accumulation of saccharides, small peptides and nucleosides, consistent even under high Na_2_SO_4_. This suggests that willow can sustain chemical energy production in leaves in the presence of sulphate, unlike with chloride, and plant-wide production of salt-tolerance supporting metabolites, underscoring the importance of understanding anion toxicity in salt tolerance and implicating organic compound sulphate incorporation in prevention of ionic imbalance. As an example, the accumulation of sulphur-containing metabolites (Fig.5i,j), including 2,3-di-O-sulfopropylglucose, aligns with the known mechanism of sugar sulphonation that promotes salt tolerance through ion regulation, osmotic tolerance, and cell wall integrity in algae (Aquino et al., 2011).

Both chloride and sulphate salts activated the terpenoid pathway (Supporting information – Fig.S9), known to mitigate salt stress in mangroves (Basyuni et al., 2009). NaCl induced significant accumulation of terpenoids from the chloroplastic methylerythritol phosphate (MEP) pathway across the whole plant, including diterpenoids as well as iridoid and secoiridoid monoterpenoids, reflecting transcriptomic changes observed in salt-tolerant poplar (Chen et al., 2018). Na_2_SO_4_ treatment similarly enriched terpenoids through the MEP pathway, but also activated the cytoplasmic mevalonate pathway in roots, also linked to salt tolerance in poplar (Wei et al., 2020). This sulphate-salt-associated activation of the mevalonate pathway increased with higher Na_2_SO_4_ levels, including enrichment of the hormone precursor abscisic aldehyde in roots and leaves. This suggests an adaptive stress response where willows are positioned to quickly synthesise ABA, signalling stomatal closure, root architecture changes, and osmoprotectant accumulation (Ryu and Cho, 2015), with ABA-aldehyde oxidase mediated regulation. In contrast, ABA precursor enrichment was not observed with NaCl, indicating chloride ion interference with signalling that may leave willows more vulnerable to salt stress.

Overall, willow responded to salt stress by modulating both core and organ-specialised metabolites (Fig.4 and Fig.7d). Within each organ, there was a tendency for greater depletion of organ-specialised compounds, over core metabolites, indicating higher resilience of core metabolism under salt stress (Fig.7d). Contrasting this, Na_2_SO_4_ treatment resulted in the enrichment of core metabolites in leaves, while NaCl enriched organ-specialised compounds. This anion-specific difference further highlights the nuanced metabolic mechanisms deployed to face distinct salt stress conditions.

## 6 Conclusion

Untargeted metabolomics uncovered extremely high diversity of metabolites in willow, the majority of which represent organ-specialised metabolites aligned with distinct physiological functions, but which also include over 250 putative plant-wide stable, or core, metabolites. The leaves had the highest metabolite diversity, but when subjected to different saline soil conditions, roots, as the frontline of exposure, had the most extensive biochemical response, mitigating ion toxicity and osmotic pressure changes to enable plant tolerance. Accompanying this was plant-wide reallocation of energy and metabolic resources, coupled with prioritisation of antioxidant flavonoids, able to protect vulnerable leaf photosynthetic functions, constituting the signature of a general willow salt response. However, the largely specialised nature of this response revealed that *in planta*, salt anion type becomes an increasingly stronger discriminating force of metabolomic profile when moving from roots to leaves. Indeed, the metabolic adjustment differed extensively in leaves, where Na_2_SO_4_ induces key components of the core metabolome to mitigate complex osmotic and toxic stress even as soil salinity increases, whereas NaCl induces more organ-specialised metabolic tools. Salinity levels in agriculture are currently categorised by soil EC, however, given the substantial anion-dependent differences observed in plant metabolic responses, soil EC measurement might be insufficient to reflect the diversity of soil salinity types, their actual osmotic potential, as well as the anion-specific plant metabolic toolkit necessary for plant salt tolerance.

## 8 Acknowledgements

We gratefully acknowledge the financial support provided from NSERC Discovery Grants (RGPIN-2017-05452 and RGPIN-2023-04863), Natural Resources Canada Forest Innovation Program Grant (CWFC1718-018 and CWFC1920-104), ECCC Environmental Damage Fund (EDF-PQ-2020b012), MITACS (IT23193), and the UCD Ad Astra Award Program. A special thank you is extended to Ariane Lafrenière, Marc-Olivier Brunette and Bruno Guerrier for their kind and effective support during harvest, and to Allan Harms from NRAL, University of Alberta, for his valuable expertise and sympathy.

## 9 Data availability statement

The data that supports the findings of this study are available in the supporting information of this article and on GNPS-MassIVE online dataset repository.

