## Supporting Information Figures for "Untargeted metabolomics reveals anion and organ-specific biochemistry of salinity tolerance in willow"

### 1 SUPPORTING INFROMATION – FIGURES

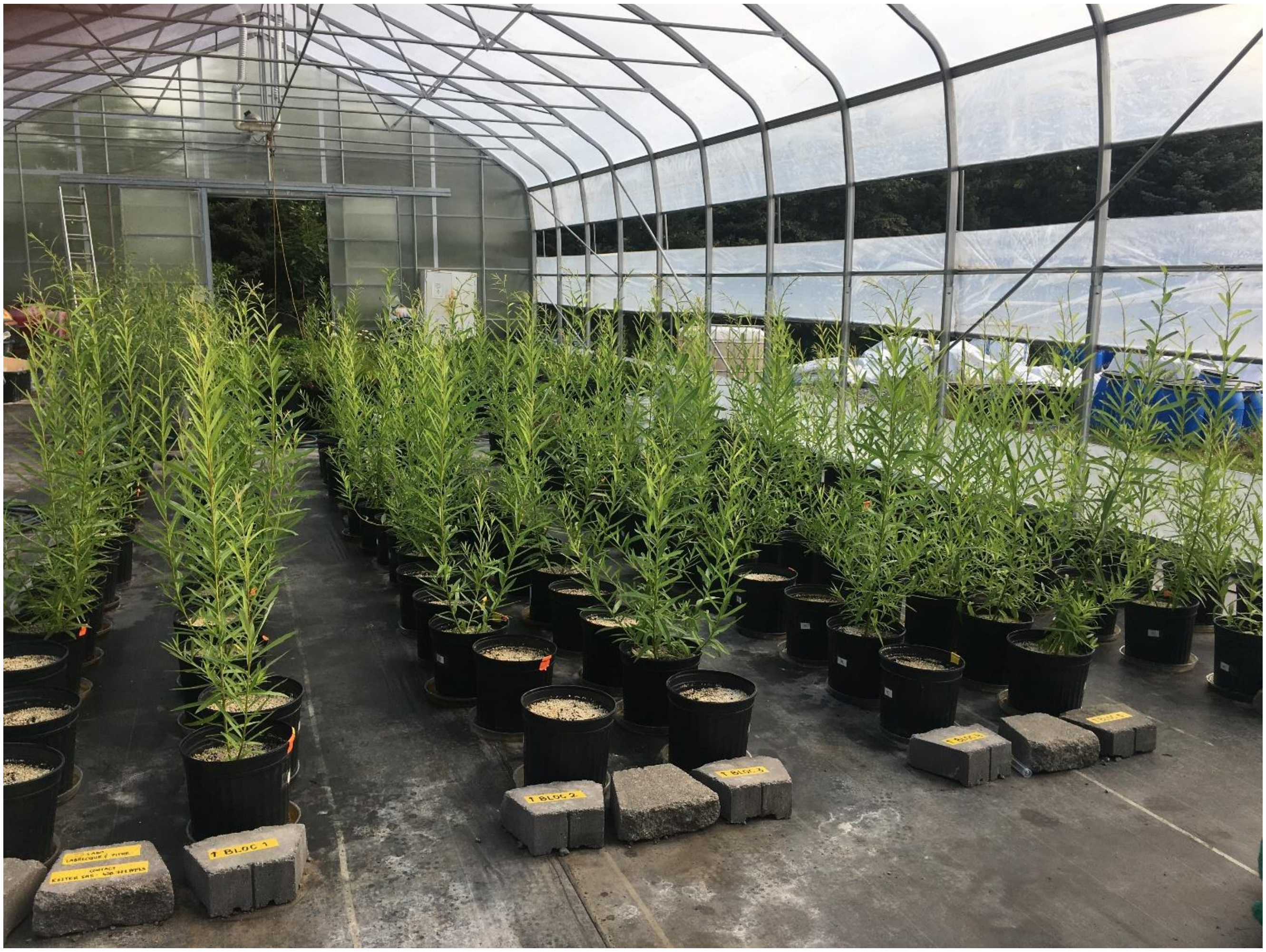


***Figure S1: Experimental setup for salt stress assessment of Salix miyabeana 'SX64'.***
Photograph showing willow plants after 67 days of growth, including a 10 days of the 28-day of treatment. Treatments involved control conditions, moderate NaCl stress, moderate $Na_2SO_4$ stress, or high $Na_2SO_4$ stress. The experiment was conducted within a polyethylene grow tunnel, with the willows cultivated in 15-liter pots.

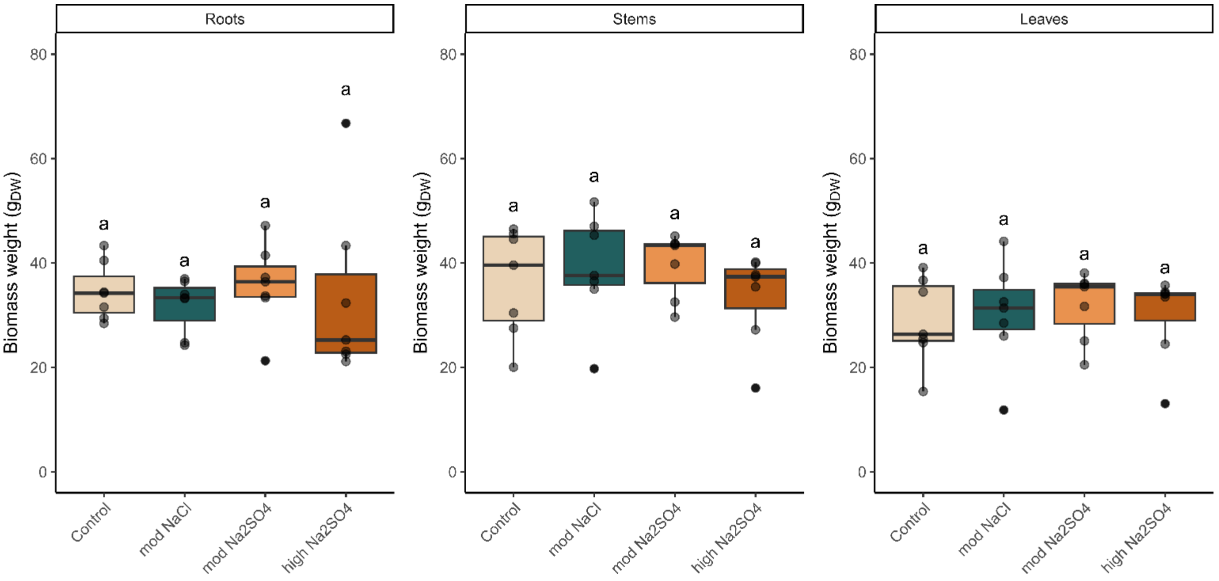


***Figure S2: Willow organs biomass (dry weight) under control and saline conditions.***
Boxes represent median and quartiles while whiskers extend to the largest value within 1.5 times the inter-quartile range (IQR) from the hinge. Dots represent individual plants (n= 7). Significant differences ($p < 0.05$) are indicated by letters were determined by two-way ANOVA followed by Tukey's post hoc test.

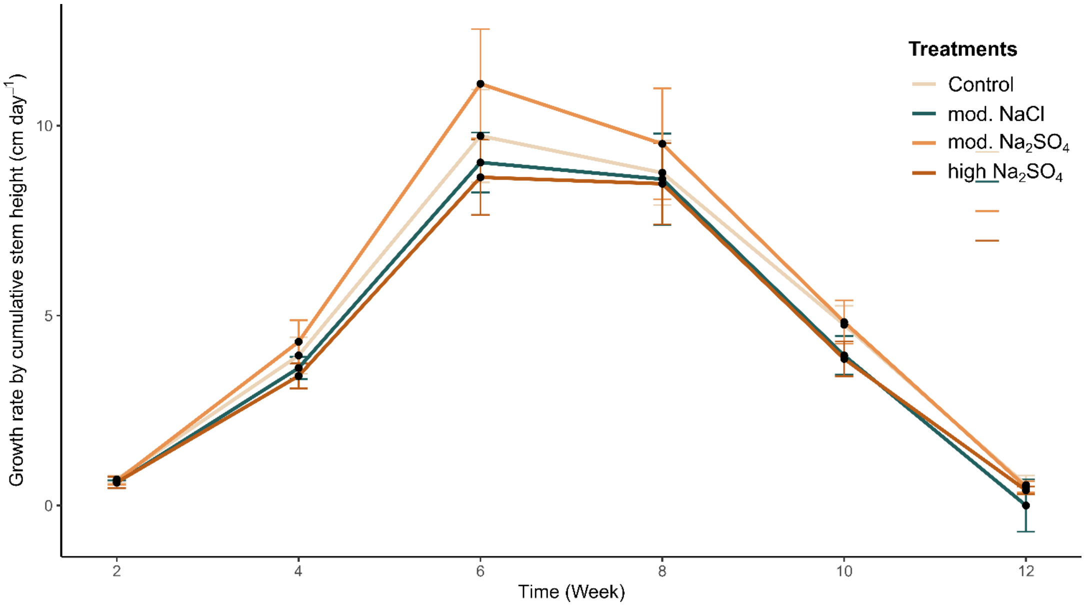


***Figure S3: Growth rate based on cumulative stem height per day.***
Stem heights were measured every 2 weeks over a 3-month experiment. Growth rate was calculated by dividing the difference in cumulative stem length between consecutive timepoints by the number of days between them (14 days). Coloured lines represent treatments: control, moderate NaCl, moderate $Na_2SO_4$, and high $Na_2SO_4$. Dots indicate mean values and error bars represent standard error (n=3). No significant differences ($p < 0.05$) were found between treatments at any timepoint, as tested by two-way ANOVA followed by Tukey's post hoc test.

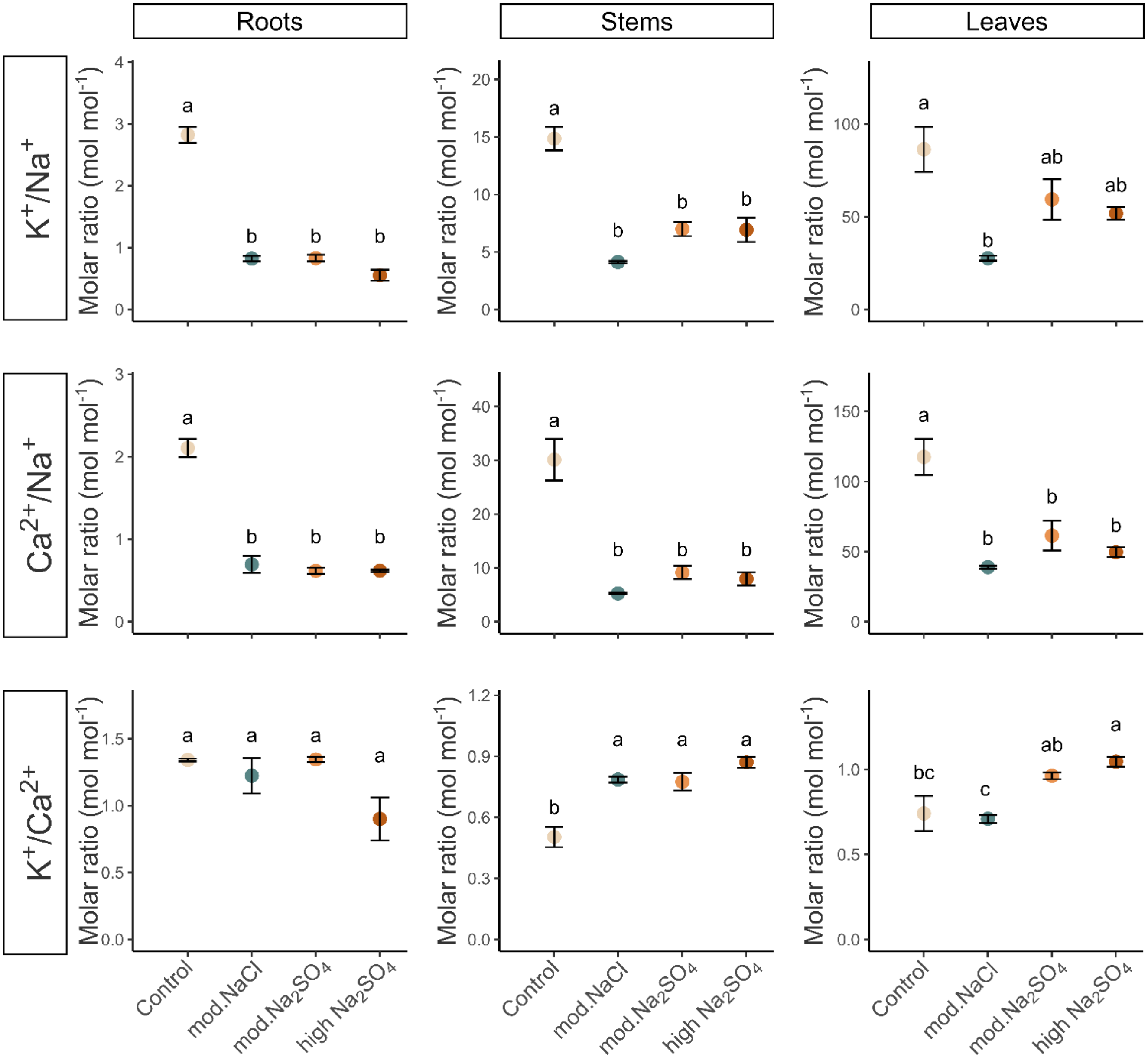


***Figure S4: Mineral elements ratios.***
Molar ratios of $K^+/Na^+$, $Ca^{2+}/Na^+$, and $K^+/Ca^{2+}$ in roots, stems and leaves of control and salt treated willows. Dots indicate mean values and error bars represent standard error (n=3). Significant differences ($p < 0.05$) are indicated by letters were determined by two-way ANOVA followed by Tukey's post hoc test.

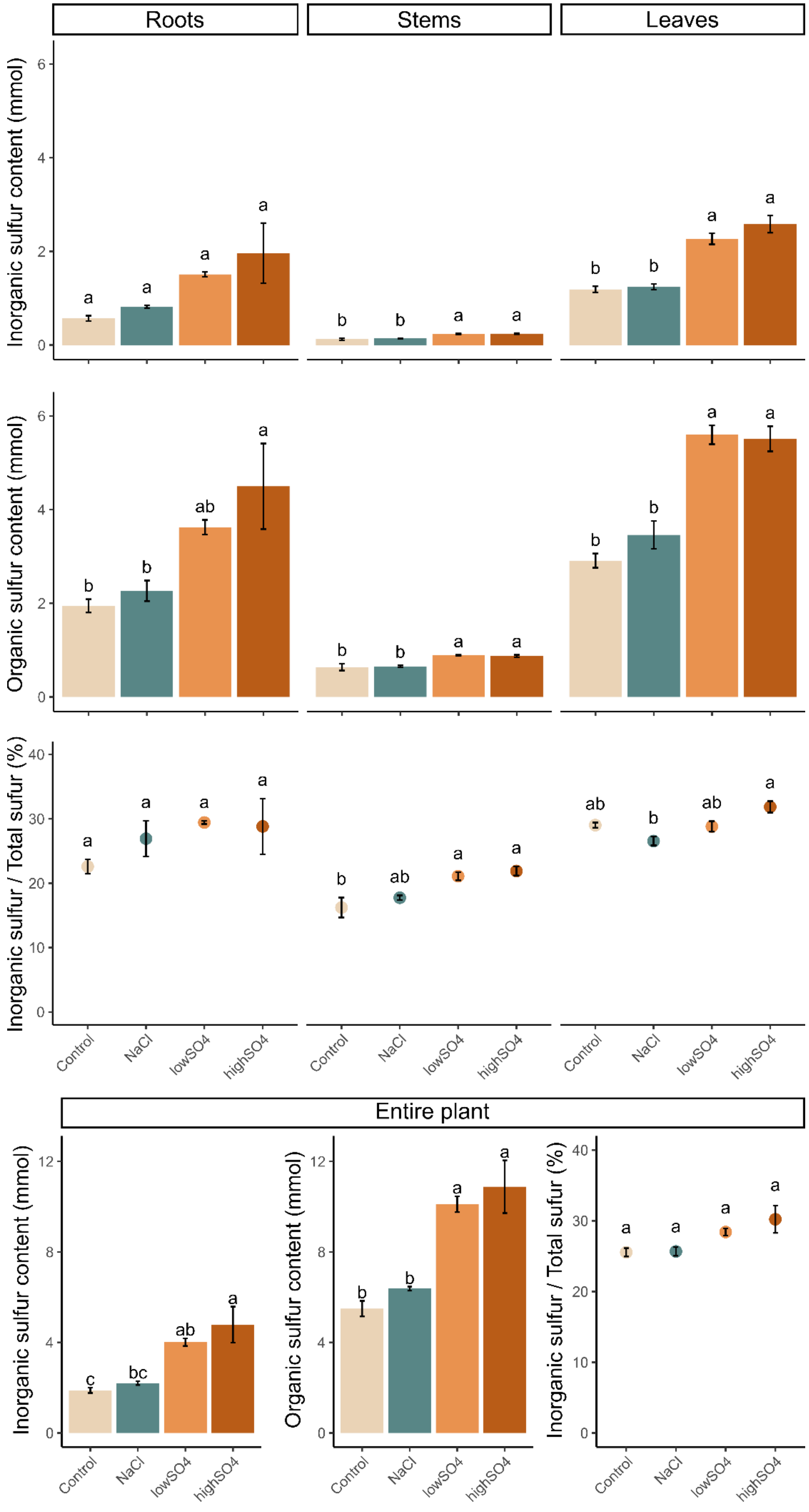


***Figure S5: Organic and inorganic sulphur.***
**(a)** inorganic sulphur ($SO_4^{2-}$) content (mmol-S), **(b)** organic sulphur content (mmol S), **(c)** proportion of inorganic sulphur based on total sulphur content (%) in roots, stems and leaves of control and salt treated willows. **(d)** Cumulative organic sulphur content, cumulative inorganic sulphur content and cumulative inorganic sulphur proportion compared to the total sulphur content in the entire plant. Bars and dots indicate mean values and error bars represent standard error (n=3).

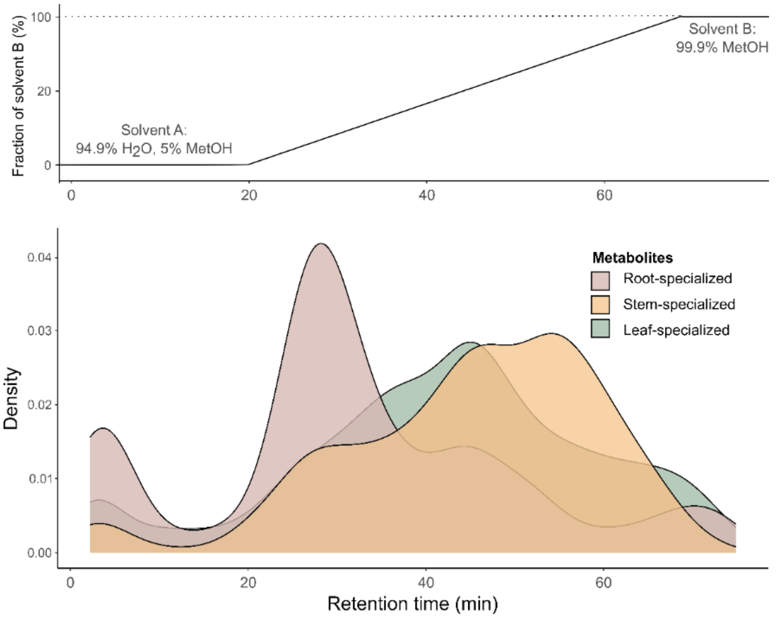


***Figure S6: Histogram of organ-specialised metabolites retention time.***
Retention time is considered as indicative of metabolite polarity. Liquid chromatography separation involved a 80-minute elution gradient: 100% A hold for 20 min (most polar fraction), then a linear increase from 0% to 100% B over 50 min and 100% B hold for 10 min (most apolar fraction).

26

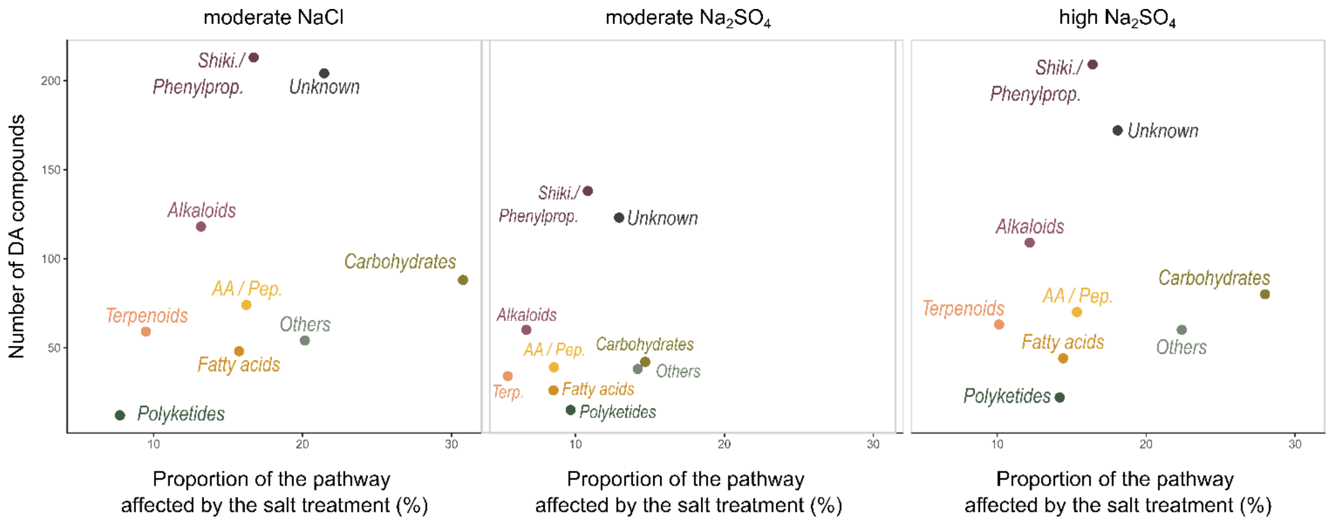


***Figure S7: Proportional change within each pathway.***
The y-axis represents the number of differentially abundant (DA) compounds within each major pathway, while the x-axis represent the proportion of these DA compounds based on the total number of compounds identified within the respective pathway.

27

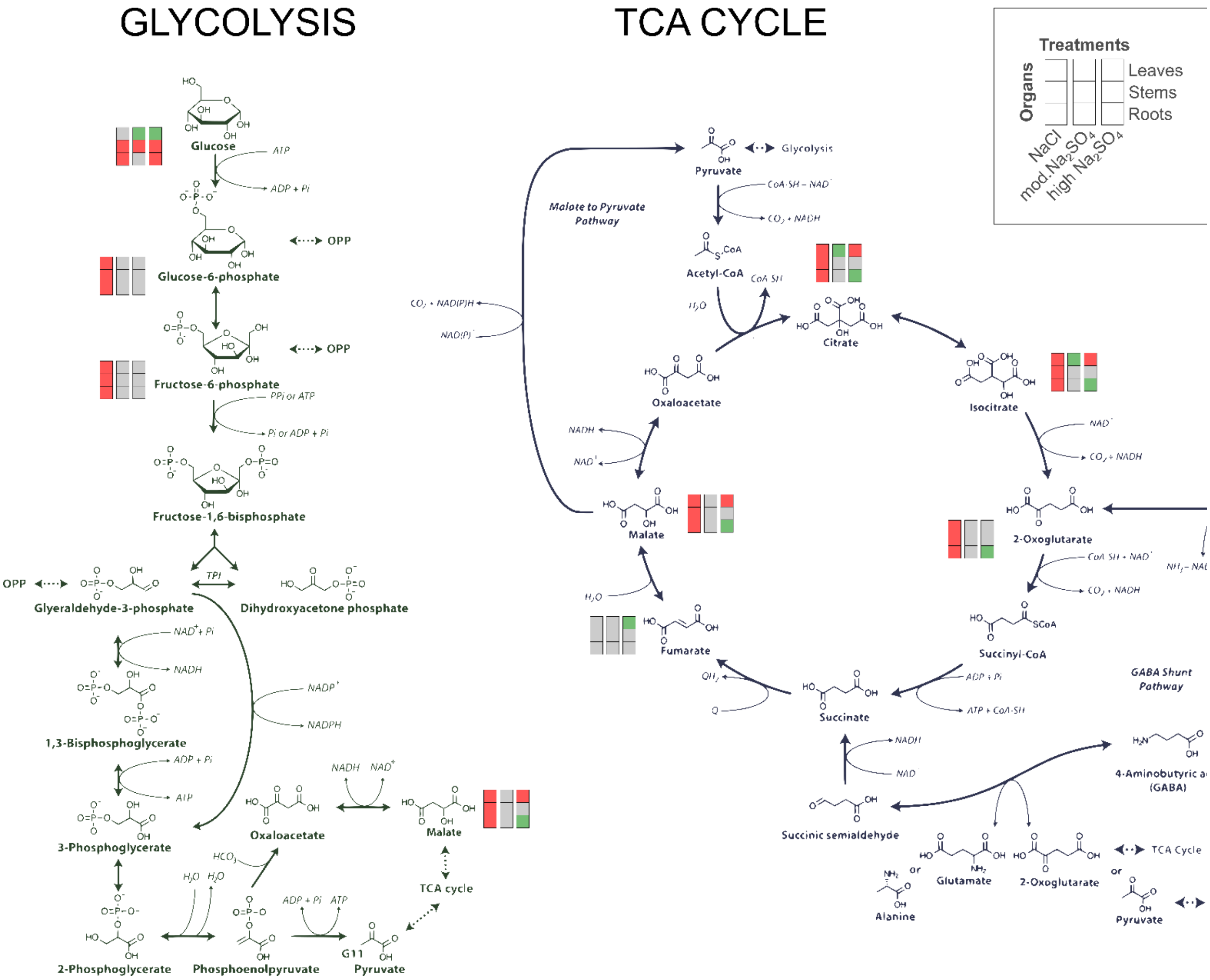


***Figure S8: Reculgation of glycolysis and TCA cycle pathways.***
Impact of salt treatment on metabolic regulation compared to control is illustrated with coloured boxes, as significantly enriched (green), significantly depleted (red) or no significantly different (grey). Identification of compounds from glycolysis and TCA cycle is based on putative annotation, without authentic standards. Significance was determine by compound chromatographic area (n=7 plants). Pathways diagram was adapted from (Bandehagh and Taylor, 2020)

28

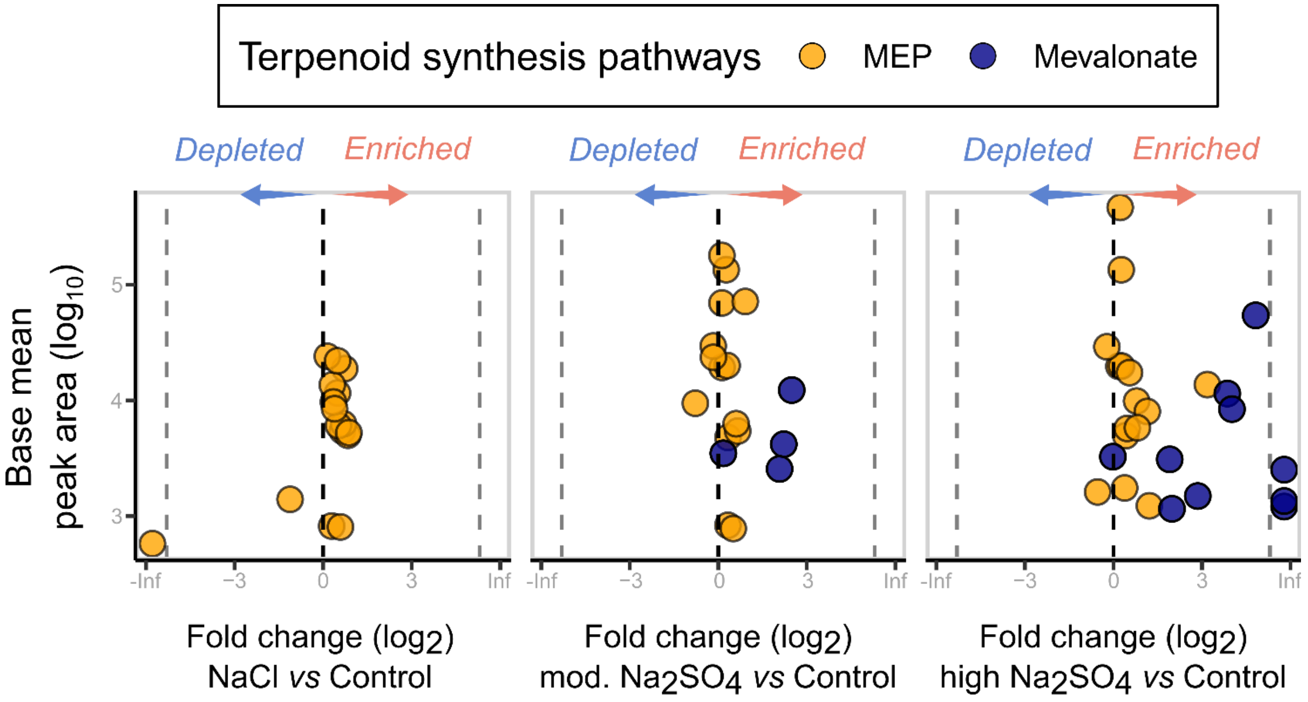


***Figure S9. Regulation of terpenoid pathway under salt stress.***

Dots are color-coded by terpenoid synthesis pathway (orange: associated to the methylerythritol phosphate pathway, blue: terpenoids associated to the mevalonate pathway). Base mean area reflects average peak area across all compared samples. Only metabolites with an MS2 spectra were included.

29
